# Astrocytic MAOB-GABA axis as a molecular brake on repair following spinal cord injury

**DOI:** 10.1101/2025.02.25.637603

**Authors:** Hye Yeong Lee, Jung Moo Lee, Hye-Lan Lee, Jiyeon Park, Heeyoung An, Eun Kyung Park, Sae Yeon Hwang, Gwang Yong Hwang, Keung Nyun Kim, Min-Ho Nam, Seung Eun Lee, Hyunji Kang, Joungha Won, Bo Ko Jang, Elijah Hwejin Lee, SunYeong Choi, Mingu Gordon Park, Sang Wook Kim, Ki Duk Park, SeungHwan Lee, C. Justin Lee, Yoon Ha

**Affiliations:** Spine & Spinal Cord Institute, Department of Neurosurgery, College of Medicine, Yonsei University, Seoul 03722, Republic of Korea; Center for Cognition and Sociality, Institute for Basic Science, Daejeon 34126, Republic of Korea; Department of Clinical Pharmacology and Therapeutics, Seoul National University College of Medicine and Hospital, Seoul 03080, Republic of Korea; NeuroBiogen Co., LTD, Seocho-gu, Seoul 06714, Republic of Korea; Brain Science Institute, Korea Institute of Science and Technology (KIST), Seoul 02792, Republic of Korea; Research Animal Resource Center, Korea Institute of Science and Technology (KIST), Seoul 02792, Korea; Center for Brain Disorders, Brain Science Institute, Korea Institute of Science and Technology (KIST), Seoul, 02792, Republic of Korea; Division of Bio-Medical Science & Technology, University of Science and Technology, Daejeon, 34113, Republic of Korea; Department of Biomedical Engineering, Ulsan National Institute of Science and Technology (UNIST), Ulsan 44919, Republic of Korea; POSTECH Biotech Center, Pohang University of Science and Technology (POSTECH), Pohang 37673, Republic of Korea

## Abstract

Neuroregeneration and remyelination rarely occur in the adult mammalian brain and spinal cord following central nervous system (CNS) injury. The glial scar has been proposed as a major contributor to this failure in the regenerative process. However, its underlying molecular and cellular mechanisms remain unclear. Here, we report that monoamine oxidase B (MAOB)-dependent excessive GABA release from reactive astrocytes suppresses CNS repair system by reducing BDNF and TrkB expression in severe spinal cord injury (SCI) animal models. Genetic deletion of MAOB in a mouse SCI model promotes both functional and tissue recovery. Notably, the selective MAOB inhibitor, KDS2010, facilitates recovery and regeneration by disinhibiting the BDNF-TrkB axis in a rat SCI model. Its dose-dependent effects were further validated in a monkey SCI model. Moreover, KDS2010 demonstrates a tolerable safety profile and dose-proportional pharmacokinetics in healthy humans during a phase 1 clinical trial. Our findings identify the astrocytic MAOB-GABA axis as a crucial molecular and cellular brake on CNS repair system following SCI and highlight translational potential of KDS2010 as a promising therapeutic candidate for SCI treatment.

## INTRODUCTION

It is firmly believed that once neurons die due to severe brain or spinal cord injury, regeneration of neurons rarely occurs in adult mammalian central nervous system (CNS)^1,2^. This failure of the mammalian CNS repair system has been attributed to the notorious glial scar, which fills in and surrounds the injured site after severe damage to the CNS^3–5^. Despite lines of evidence, how glial scars establish a brake on the CNS repair system has remained a mystery for several decades.

The role of glial scar in injured mammalian tissue is complicated^6^. Some studies reported a neuroprotective role of reactive astrocytes in mild acute injury conditions^4,7^. In contrast, numerous studies have suggested that reactive astrocytes inhibit neuroregeneration and tissue recovery by releasing inhibitory molecules such as cytokines and chondroitin sulfate proteoglycan (CSPG), which are proposed to serve as a barrier to CNS axon extension^8–10^. Although there appear to be sufficient lines of circumstantial evidence supporting the idea that scar-forming reactive astrocytes play a detrimental role in neuroregeneration and recovery, the debate continues. This ongoing debate is probably due to the lack of the precise molecular identity acting as a brake on the CNS repair system and the absence of molecular and pharmacological tools to reduce glial scars effectively.

We have recently developed powerful molecular and pharmacological tools to abrogate reactive astrogliosis and glial scar in various neuroinflammatory conditions^11–16^. These tools can effectively block GABA- and H_2_O_2_-production by targeting either the putrescine degradation pathway via MAOB or the putrescine synthesis pathway via the urea cycle. Notably, the reversible and selective MAOB inhibitor, KDS2010, has reversed learning and memory impairment in Alzheimer’s disease model^11,12^, motor impairment in Parkinson’s disease model^14^, and motor impairment in white matter stroke model^13^. These findings highlight the inhibitory roles of MAOB-dependent astrocytic GABA via the putrescine degradation pathway in multiple pathological conditions. However, the potential inhibitory role of astrocytic GABA in recovery after SCI has not been investigated.

This study identifies that MAOB-dependent excessive GABA from the scar-forming reactive astrocytes acts as a molecular and cellular brake on recovery after SCI by reducing BDNF and TrkB expression. Disinhibiting the BDNF-TrkB axis through genetic or pharmacological inhibition of the MAOB-GABA axis promotes recovery in severe SCI animal models, from rodent to non-human primate. Additionally, findings from the phase 1 clinical trial of KDS2010 demonstrate its safety and tolerability in human subjects. The integration of preclinical and clinical data provides strong support for the therapeutic potential of MAOB inhibition in SCI recovery, offering new avenues for treatment.

## RESULTS

### Astrocytic MAOB impedes functional and tissue recovery in mouse SCI model

To assess the pathological role of MAOB in SCI, we examined the recovery after SCI using three types of genetically modified mouse line for *Maob*: MAOB knockout (KO), astrocyte-specific MAOB conditional knockout (aKO), and astrocyte-specific MAOB conditional overexpression (aOE). The severe crush injury was carried out at the thoracic vertebra number 10 in each mouse line. After generating severe mouse SCI model, we assessed the Basso mouse scale (BMS) for locomotion score^17^ to test the functional recovery once a week for total of 11 weeks (1 week before and 10 weeks after the surgical operation) (Fig. 1a). We found that MAOB KO with SCI group (MAOB KO+SCI) showed a significant recovery of motor function compared to WT with SCI group (WT+SCI) from post-injury (PI) 3 week (3w), whereas sham-operated WT or MAOB KO group showed no sign of motor dysfunction (Fig. 1b). Notably, at PI 10w, MAOB KO+SCI achieved a score of 3.11±0.22 (N=35), while WT+SCI scored only 1.62±0.16 (N=34) (Fig. 1b). According to the BMS scoring criteria^17^, a score of 1∼2 is defined as slight or extensive ankle movement and a score of 3 indicates the threshold at which the animal can support its own body weight^17,18^. Therefore, the difference between a score of 1∼2 and 3 in a severe mouse SCI model indicates a substantial functional recovery. Similarly, aKO+SCI also showed a significant functional recovery in motor function from PI 2w compared to CTL+SCI (Fig. 1c). In contrast, aOE+SCI exhibited more severe motor dysfunction than CTL+SCI (Fig. 1d). These results indicate that MAOB, specifically expressed in astrocyte, hinders functional recovery after SCI.

**Fig. 1.**
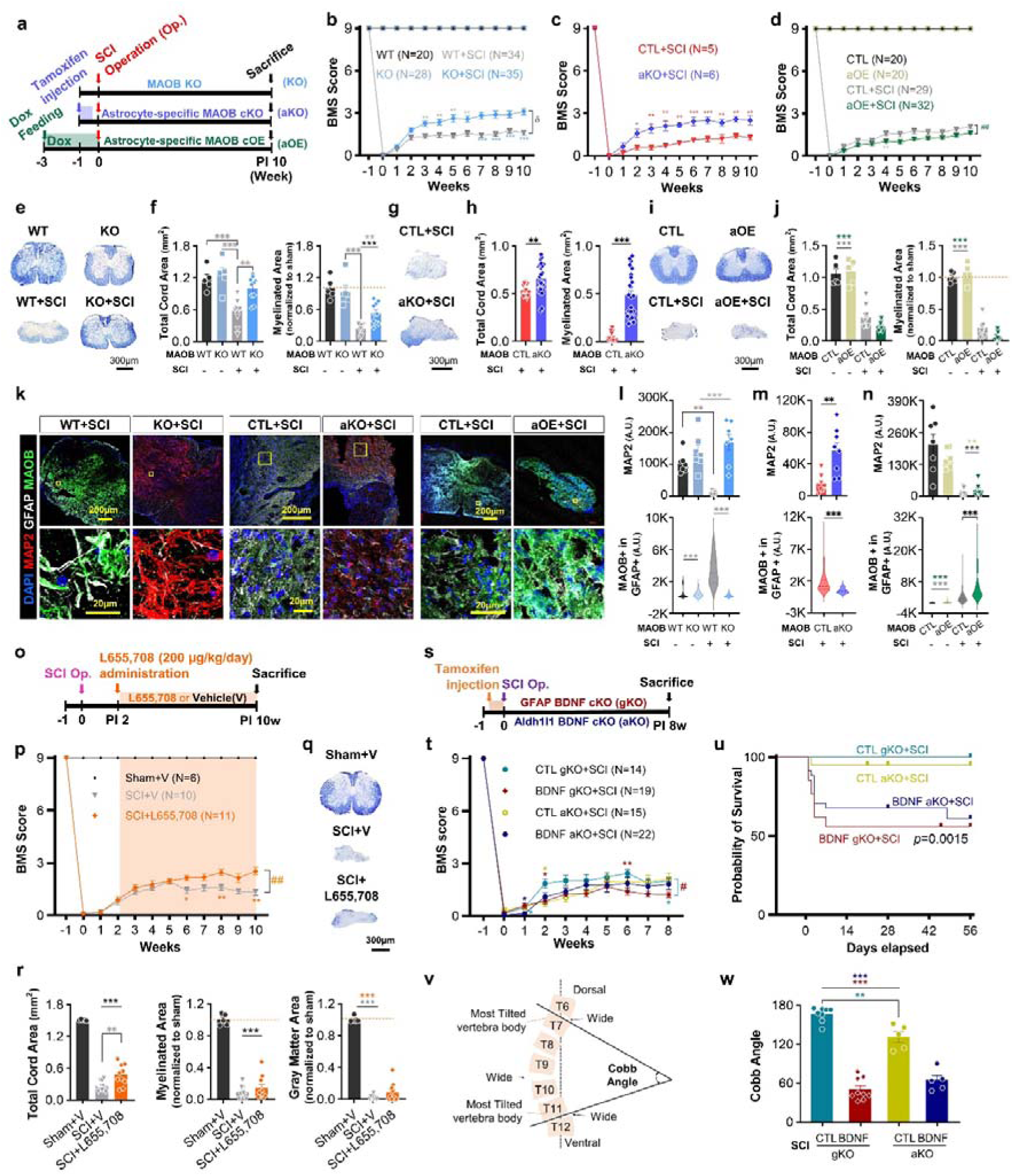
Astrocytic MAOB and GABA impedes functional and tissue recovery, while astrocytic BDNF is critical for survival after SCI. **a** Experimental timelines using MAOB KO, aKO, and aOE. **b** BMS score of each group (MAOB WT, WT+SCI, MAOB KO, and KO+SCI) for a total of 11 weeks (1 week before and 10 weeks after the surgical operation). **c, d** BMS score of each group for aKO (**c**), and aOE (**d**) for a total of 11 weeks. **e, g, i** EC staining of spinal cord tissues in each group for MAOB KO (**e**), aKO (**g**), and aOE (**i**) at PI 10w. **f, h, j** Total cord area (left) and myelinated area (right) of each group for MAOB KO (**f**), aKO (**h**), and aOE (**j**) in EC staining. Myelinated area was normalized to WT in group for MAOB KO and aOE. **k** Confocal images of the injured area in each group for MAOB KO (left), aKO (middle), and aOE (right) stained with anti-MAP2 (red), GFAP (white), MAOB (green) antibodies, and DAPI (blue) at PI 10w. Each yellow box in merged images indicates the magnified region of interest. **l, m, n** Mean intensity of the MAP2 (top) and GFAP-positive MAOB (bottom) in each group for MAOB KO (**l**), aKO (**m**), aOE (**n**) at PI 10w. **o** Experimental timeline using 8-week-old WT mice with the SCI operation and inverse agonist for α5-containing GABA_A_ receptor, L655,708. **p** BMS score of each group (Sham+V, Sham+L655,708, SCI+V, and SCI+ L655,708) for a total of 11 weeks. Orange shade indicates the duration of L655,708 administration. **q** EC staining of spinal cord tissues in each group at PI 10w. **r** Total cord area (left), myelinated area (middle), and grey matter area (right) of each group in EC staining. **s** Experimental timeline using BDNF gKO and aKO. **t** BMS score of each group for a total of 9 weeks. **u** Survival curves of each group. **v** Schematic figure showing the calculation method for cobb angle. **w** Calculated cobb angle of each group at PI 8w (left) and representative image showing spine deformity of BDNF gKO+SCI (right). All data are expressed as mean ± S.E.M. \**P* < 0.05; \*\**P* < 0.01; \*\*\**P* < 0.001; n.s., not significant. #, ##, ###, and δ indicate *P* < 0.05, *P* < 0.01, *P* < 0.001, and *P* < 0.0001 for linear mixed model.

We next examined the extent of injury at the tissue level by performing the Eriochrome Cyanine (EC) staining at PI 10w. Compared to WT and MAOB KO, WT+SCI showed significantly reduced total spinal cord, myelinated, and grey matter areas, which were all significantly recovered in MAOB KO+SCI (Fig. 1e, f). Similarly, aKO+SCI showed significantly larger total spinal cord, myelinated, and grey matter areas than CTL+SCI (Fig. 1g, h), while aOE+SCI showed a tendency toward smaller total spinal cord, myelinated, and grey matter areas than CTL+SCI at PI 10w (Fig. 1i, j). These results indicate that astrocytic MAOB impedes functional and tissue recovery after SCI.

To test whether this functional/tissue recovery correlates with signs of neuroregeneration and astrocyte reactivity, we performed immunohistochemistry with antibodies against MAP2 (neuronal marker), GFAP (astrocytic marker), and MAOB (reactive astrocytic marker)^11^ in the ventral region of the spinal cord. The intensity of MAP2 was significantly decreased in WT+SCI, whereas it was fully recovered in MAOB KO+SCI (Fig. 1k, l and Supplementary Fig. 1a), indicating a positive correlation between signs of neuroregeneration and functional/tissue recovery. On the other hand, the intensity of astrocytic GFAP and MAOB of WT+SCI were significantly higher than that of WT, MAOB KO, or MAOB KO+SCI (Fig. 1k, l and Supplementary Fig. 1a), indicating a negative correlation between astrocyte reactivity and functional/tissue recovery. Additionally, we observed a significant increase in MAP2 intensity and a marked reduction of astrocytic GFAP and MAOB in aKO+SCI compared to CTL+SCI (Fig. 1k, m and Supplementary Fig. 1b). In contrast, aOE+SCI showed no recovery in MAP2 intensity and a significant elevation in astrocytic GFAP and MAOB compared to CTL+SCI (Fig. 1k, n and Supplementary Fig. 1c). These results further confirmed the positive correlation between neuroregeneration and recovery, as well as the negative correlation between astrocyte reactivity and recovery.

### MAOB-dependent GABA is the brake on functional and tissue recovery after SCI

To investigate the molecular and cellular identity of MAOB-dependent brake in SCI, we considered the observed negative correlation between the astrocyte reactivity and the functional/tissue recovery after SCI, which is reminiscent of a proposed concept of a reciprocal relationship between MAOB-dependent GABA and proBDNF expression in hypertrophic astrocytes^19^. We have previously defined two types of hypertrophic hippocampal astrocytes into either GABA-positive/proBDNF (pro-form of brain-derived neurotrophic factor)-negative ‘reactive astrocytes’ or proBDNF-positive/GABA-negative ‘active astrocytes’ in response to either aversive or beneficial environmental stimuli, respectively^19^. This definition led us to investigate the roles of GABA and proBDNF in SCI.

Tonically released augmented GABA from reactive astrocytes can bind to the extrasynaptic GABA_A_ receptors, primarily containing the α_5_ subunit^20^, resulting in the inhibition of neighboring neurons^11^. Therefore, we examined whether the tonic GABA inhibition in SCI contributes to the failure of function/tissue recovery. We assessed the BMS for locomotion score in the severe mouse SCI model with intraperitoneal (i.p.) injection of L655,708, an inverse agonist for α5-containing GABA_A_ receptor, from the sub-acute phase of SCI (PI 2w), when the substantial amount of glial scar is present^21^ (Fig. 1o). We found that SCI+L655,708 showed a significant functional improvement compared to SCI+V from PI 6w, whereas Sham+V or Sham+L655,708 showed no sign of motor dysfunction (Fig. 1p). We then examined the extent of injury at the tissue level by performing the EC staining at PI 10w. Compared to SCI+V, SCI+L655,708 showed significantly increased total spinal cord and had a tendency toward bigger myelinated and grey matter areas (Fig. 1q, r). We confirmed the positive correlation between signs of neuroregeneration and functional/tissue recovery by observing an increasing tendency of MAP2 intensity in SCI+L655,708 compared to SCI+V (Supplementary Fig. 2a, b). Because the GABA receptor is downstream of MAOB-dependent GABA production and release from reactive astrocytes, SCI+L655,708 showed no change of astrocyte reactivity, as indicated by the intensity of astrocytic GFAP and MAOB, compared to SCI+V (Supplementary Fig. 2a, b). These results suggest that astrocytic MAOB-dependent GABA is the molecular and cellular brake on the repair process in SCI.

Previous studies have demonstrated that oxidative stress, such as reactive oxygen species (ROS), facilitates neurodegeneration after SCI^22–24^. Moreover, we recently reported that ROS, especially H_2_O_2_, is generated through the MAOB-dependent putrescine degradation pathway in severe reactive astrocytes^25^. Thus, we tested the effect of ROS scavenging on recovery after SCI by administering the ROS scavenger, AAD2004, via i.p. injection starting from PI 2w. However, it did not impact functional/tissue recovery and astrocyte reactivity (Supplementary Fig. 2c-h). These results are consistent with the previous report demonstrating the effectiveness of antioxidant therapy only in the acute phase, not in the sub-acute or chronic phase, of SCI^22^.

### Astrocytic BDNF is critical for survival in mouse SCI model

Next, we examined the role of astrocytic BDNF in SCI using two types of astrocyte-specific BDNF cKO mice. These mice were generated by crossing GFAP-CreER^T2^ or Aldh1l1-Cre/ER^T2^ with *Bdnf* floxed mice and subsequently injecting the tamoxifen for five consecutive days from 7-week-old (BDNF gKO and BDNF aKO) (Fig. 1s). We observed that tamoxifen-treated mice (BDNF gKO+SCI and BDNF aKO+SCI) showed more severe motor dysfunction and lower survival rate than sunflower oil-treated control mice (CTL gKO+SCI and CTL aKO+SCI) (Fig. 1t, u). Although total cord, myelinated, and grey matter areas did not show any difference (Supplementary Fig. 3a, b), tamoxifen-treated mice showed a more severe cobb angle of the spine, indicating the extent of spine deformities, compared to sunflower oil-treated mice (Fig. 1v, w). Interestingly, there were no significant differences in MAOB, GFAP, and astrocytic MAOB levels between the two groups (Supplementary Fig. 3c, d), suggesting that BDNF signaling is downstream of astrocyte reactivity. Moreover, tamoxifen-treated mice showed no change in GABA levels, accompanied by significantly lower proBDNF levels compared to sunflower oil-treated mice (Supplementary Fig. 3e, f). These results provide novel mechanistic insights, positioning astrocytic GABA as an upstream regulator of BDNF signaling and astrocytic proBDNF as a critical contributor to survival after SCI.

### KDS2010, a selective MAOB inhibitor, causes functional and tissue recovery in rat SCI model

Next, we tested whether pharmacological inhibition of MAOB facilitates recovery after SCI. To test this, we used the selective and reversible MAOB inhibitor KDS2010, which circumvents the shortcomings of irreversible MAOB inhibitors^12^. We first evaluated the effect of KDS2010 on recovery after SCI at a dose of 10mg/kg (mpk), previously reported to be effective in treating other neurological diseases^12–14^. KDS2010 was administered by drinking *ad libitum* to both sham-operated and SCI-operated groups, starting from the sub-acute phase (PI 2w) (Fig. 2a). We substituted the mouse SCI model with the rat SCI model, which is reported to be more optimal for pharmacological evaluation and closer to human SCI^26^. We performed the Basso, Beattie and Bresnahan (BBB) locomotor test^27^ to assess the functional recovery once a week for a total of 11 weeks (Fig. 2a). As a result, SCI with KDS2010 treatment group (SCI+KDS 2w) showed a significant motor recovery compared to SCI with vehicle group (SCI+V) starting from PI 3w, reaching an average BBB score of 10.62 at PI 10w, enabling walking with weight on the sole (Fig. 2b, c and Supplementary Movie 1). In addition to BBB locomotor test, we performed the ladder rung test by measuring the percentage of hindlimb steps without slipping as rats walked along a horizontal ladder^28^ (Fig. 2d, e). We observed significant motor recovery in SCI+KDS 2w, reaching about 65% compared to sham-operated rats, while the SCI+V showed no signs of recovery (Fig. 2d and Supplementary Movie 2). We then assessed the tissue recovery using EC staining on cross and longitudinal sections of the spinal cords. Compared to Sham+V and Sham+KDS, SCI+V showed significantly reduced total spinal cord, myelinated, and grey matter, along with an enlarged cavity size (Supplementary Fig. 4a, b). These pathological changes were all significantly recovered in SCI+KDS 2w (Supplementary Fig. 4a, b). These results indicate that MAOB inhibition with KDS2010 facilitates functional/tissue recovery after SCI.

**Fig. 2.**
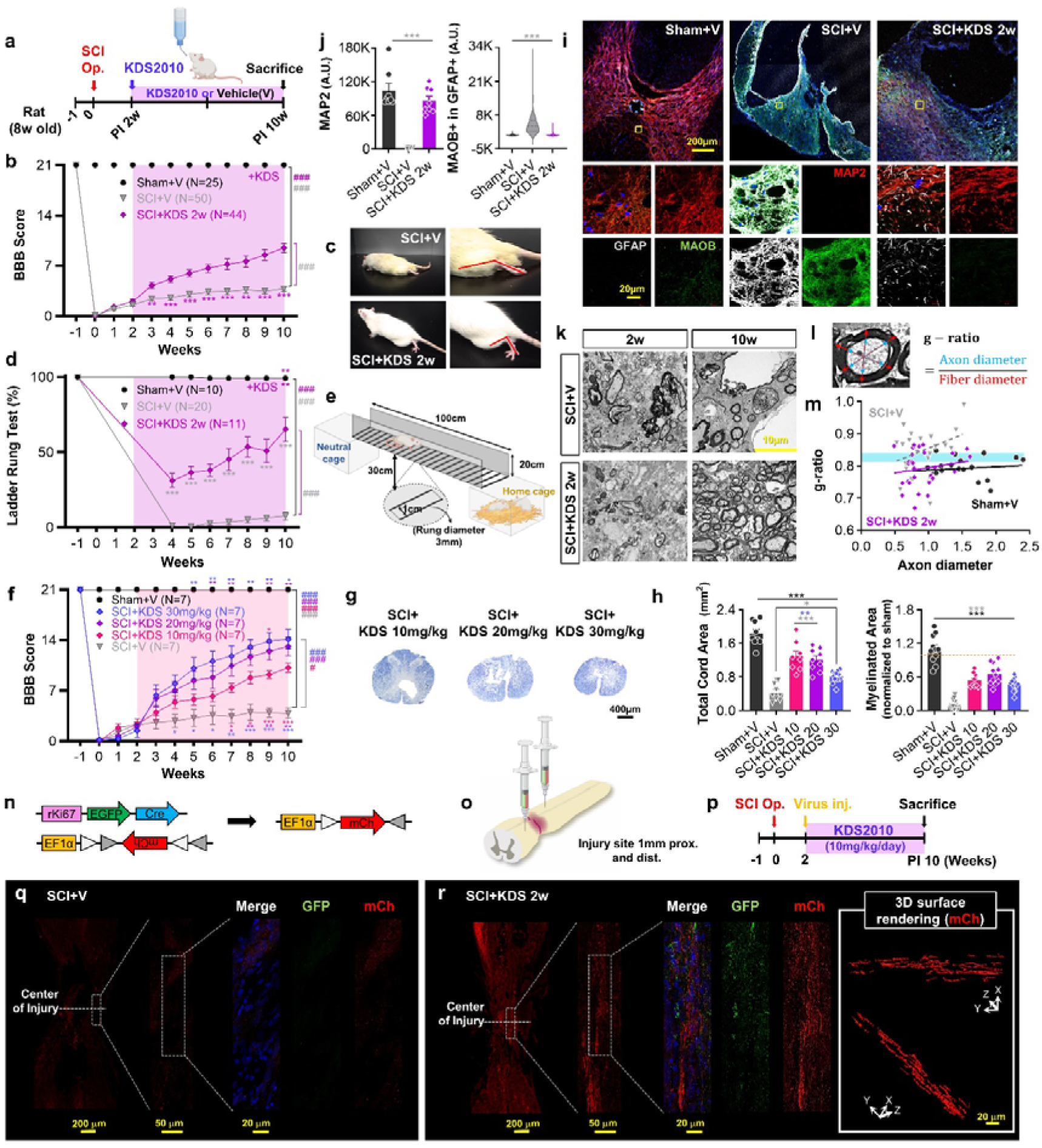
MAOB inhibition with KDS2010 causes functional/tissue recovery and neuroregeneration. **a** Experimental timelines using 8-week-old rats with the SCI operation and treatment with the MAOB inhibitor, KDS2010, from 2 weeks after (sub-acute) SCI. **b** BBB score of each group (Sham+V, SCI+V, SCI+KDS 2w) for a total of 11 weeks (1 week before and 10 weeks after the surgical operation). **c** Hindlimb movement observation in the open field test for SCI+V and SCI+KDS 2w. **d** Percentage of hindlimb steps without slipping in each group with sub-acute phase KDS2010 treatment. **e** Schematic figure of the ladder rung test apparatus. Created with BioRender.com. **f** BBB score of each group with three different doses of KDS2010 (Sham+V, SCI+V, SCI+KDS 10mpk, 20mpk, and 30mpk) for a total of 11 weeks. All doses of KDS2010 were treated from the sub-acute phase. **g** EC staining of spinal cord tissues in each dose at PI 10w. **h** Total cord area and myelinated area of each group in EC staining. **I** Confocal images of the injured area in each group stained with anti-MAP2 (red), GFAP (white), and MAOB (green) antibodies at PI 10w. Each yellow box in merged images indicates the magnified region of interest. **j** Mean intensity of the MAP2 (left) and GFAP-positive MAOB (right) in each group. **k** TEM images of spinal cord tissues at PI 2w and 10w in SCI+V and SCI+KDS 2w. **l** Calculated method for g-ratio. **m** Calculated g-ratio in each group. Blue shade indicates the optimal range of g-ratio (0.790 ± 0.005). **n** Schematic figure showing DNA constructs in each virus and the resultant of recombination. **o** Schematic figure showing the virus injection sites at 1 mm proximal and distal the injured area. **p** Experimental timeline using rats with the SCI operation, virus injection, and sub-acute phase treatment of KDS2010 at 10 mpk. **q** Confocal images of the injured area in SCI+V stained with anti-GFP (green), mCh (red) antibodies and DAPI (blue) at PI 10w. **r** Confocal images of the injured area (left) and 3D surface rendering images of mCh signals in SCI+KDS 2w at PI 10w. Shaded areas in **b**, **d, k** indicate the duration of KDS2010 administration.

Next, we explored the dose-dependent effect of KDS2010 on recovery. The results showed a dose-dependent improvement in the BBB locomotor scores with 10, 20, and 30mpk at PI 10w (Fig. 2f), accompanied by a significant tissue recovery (Fig. 2g, h). Furthermore, we evaluated the effects of KDS2010, at the sub-saturating concentration of 10mpk, across different phases of SCI on recovery. Following MAOB inhibition from the acute phase (PI 1d), the animals were divided into two groups: one with a significant recovery and the other with no recovery (Supplementary Fig. 7a, b). In the BBB locomotor test, we found that SCI+KDS 1d/Re achieved a stable gait with an average BBB score of 15.36 at PI 10w, while SCI+KDS1d/No did not show any behavioral recovery (Supplementary Fig. 7b), suggesting a potential association with the activation of an alternative pathway, such as DAO, as previously reported^29^. Indeed, the alternative DAO pathway was activated in SCI+KDS 1d/No (Supplementary Fig. 7l, m). At the tissue level, SCI+V and SCI+KDS 1d/No showed significantly reduced total spinal cord and myelinated areas, which were all significantly recovered in SCI+KDS 1d/Re at PI 10w (Supplementary Fig. 7c, d). When we inhibited MAOB from the chronic phase (PI 6w), SCI+KDS 6w also showed a significant recovery in both BBB locomotor test and ladder rung test (Supplementary Fig. 8a-c), accompanied by a tendency toward larger total spinal cord and myelinated areas than SCI+V (Supplementary Fig. 8d, e). However, the extent of recovery was less than that observed in SCI+KDS 1d/Re and SCI+KDS 2w. Although the recovery group with acute phase treatment (SCI+KDS 1d/Re) showed better recovery than the group with sub-acute phase treatment (SCI+KDS 2w), the acute phase treatment appeared more variable due to the division into two groups with high recovery and no recovery. These results suggest that KDS2010 treatment from the sub-acute phase induces consistent and less variable functional and tissue recovery after SCI.

### KDS2010 induces signs of neuroregeneration and remyelination in rat SCI model

Next, we examined signs of neuroregeneration and the extent of astrocyte reactivity upon pharmacological inhibition of MAOB using KDS2010. Immunohistological analysis showed that MAP2 intensity was significantly reduced in SCI+V, whereas it was fully restored in SCI+KDS 2w (Fig. 2i, j). The expression of astrocytic GFAP and MAOB was aberrantly increased in SCI+V, whereas it returned to normal level in SCI+KDS 2w (Fig. 2i, j). Moreover, immunostaining with antibodies against Tuj1 and GFAP in longitudinal sections showed consistent findings (Supplementary Fig. 5a). Similar results were observed with doses of 20mpk and 30mpk (Supplementary Fig. 6a, b) and with treatment from acute (Supplementary Fig. 7e-i) or chronic phase (Supplementary Fig. 8f, g). Again, these results indicate a positive correlation between signs of neuroregeneration and functional/tissue recovery, as well as a negative correlation between astrocyte reactivity and functional/tissue recovery.

It has been previously reported that functional behavioral recovery is associated not only with neuroregeneration but also with remyelination^30,31^. To assess the impact of MAOB inhibition on the extent of myelination after SCI, we performed transmission electron microscopy (TEM) at PI 2, 4, 6, and 10w. The TEM images revealed that the severe loss of myelination, which was observed throughout the PI 10w period in SCI+V, gradually increased and fully recovered in SCI+KDS 2w at PI 10w (Fig. 2k and Supplementary Fig. 4e, f). Microstructures in TEM images also showed the evidence for the remyelination (Supplementary Fig. 4c). Moreover, we observed that the g-ratio, a functional and structural index of axonal myelination or demyelination^32^, in SCI+KDS 2w (0.792 ± 0.010) was fully restored to the optimal range of 0.790 ± 0.005^32^ (Fig. 2l, m and Supplementary Fig. 4f). In addition, we confirmed a gradual increase in the number of myelinated axons and a gradual decrease in the density of cavities in SCI+KDS 2w by using the Toluidine Blue (TB) staining (Supplementary Fig. 4g, h). Finally, immunohistochemistry with antibodies against proliferation marker Ki67 and MBP (myelin basic protein; oligodendrocyte marker) at PI 10w showed a substantial population of oligodendrocytes positive for both Ki67 and MBP in SCI+KDS 2w (Supplementary Fig. 5d, e). These findings indicate that MAOB inhibition by KDS2010 facilitates oligodendrocyte proliferation and remyelination after SCI.

### KDS2010 facilitates neuronal proliferation and axon regeneration in rat SCI model

There is a possibility that the increase in MAP2 and Tuj1 expression upon MAOB inhibition might be solely a result of a protective effect or invading axons from the peripheral nervous system rather than regenerating axons from the proliferated neurons. Thus, we tested the existence of proliferating neurons at the injury site using immunohistochemistry with antibodies against Ki67 and NeuN at PI 10w. As a result, we found a significant increase in neurons positive for both Ki67 and NeuN compared to SCI+V (Supplementary Fig. 5b, c). We then tested whether axon fibers from the proliferating neurons can traverse the injury center upon MAOB inhibition. We employed a dual viral strategy using a newly developed Ki67 promoter, which consists of a proliferation-dependent virus expressing Cre recombinase tagged with EGFP and a Cre-dependent virus expressing fluorescent protein mCherry (mCh) (Fig. 2n). This strategy enables permanent labelling of both proliferating and proliferated cells with mCh from the time of virus injection (PI 2w) (Fig. 2o, p). This approach was validated in the dentate gyrus of rats, where the percentage of proliferating and proliferated cells was 5.14±0.53% (n=3) for 2 weeks, which aligns with a previous study^33^ (Supplementary Fig. 5f, g). In SCI+KDS 2w, we observed a small number of EGFP-positive signals and many mCh-positive signals, while SCI+V showed almost no fluorescence signals (Fig. 2q, r). Remarkably, mCh-positive signals in SCI+KDS 2w traversed the core of the injury site and can be predicted to be axon fibers by 3-dimensional (3D) rendering (Fig. 2r). These results indicate that pharmacological inhibition of MAOB using KDS2010 allows axon fibers from the proliferating and proliferated neurons to traverse through the injury site.

### KDS2010 reduces astrocytic GABA levels and enhances proBDNF and TrkB expression in rat SCI model

We suggested novel mechanistic insight by identifying astrocytic GABA as an upstream regulator of BDNF signaling (Fig. 1o-w). Additionally, numerous reports have demonstrated that BDNF-TrkB signaling is critically involved in neuroregeneration and functional recovery after SCI^34,35^. Based on this evidence, we further investigated whether MAOB inhibition with KDS2010 reduces GABA levels and enhances BDNF-TrkB signaling, thereby promoting neuroregeneration and functional/tissue recovery in a severe rat SCI model. We first assessed the cellular content of GABA in GFAP-positive astrocytes using immunohistochemistry, following treatment with 10mpk of KDS2010 from the sub-acute phase. The intensity of GABA, along with GFAP, significantly increased in SCI+V compared to Sham+V or Sham+KDS, while it significantly recovered by MAOB inhibition in SCI+KDS 2w (Supplementary Fig. 9a-c), even at different doses and phases of KDS2010 treatment (Supplementary Fig. 6e, f, Supplementary Fig. 7l, m, Supplementary Fig. 8h-k).

To see if the astrocytic GABA can be released and contribute to tonic GABA inhibition in neighbouring neurons, we measured GABA_A_ receptor-mediated currents from the lamina II dorsal horn neurons at PI 10w (Fig. 3a). We observed that tonic GABA current density was significantly increased in SCI+V compared to Sham+V, whereas this aberrant tonic GABA current was significantly recovered in SCI+KDS 2w (Fig. 3b, c). We found no alteration in the extrasynaptic GABA_A_ receptor expression and in the frequency and amplitude of spontaneous inhibitory postsynaptic currents (sIPSCs) (Fig. 3c). Furthermore, in the ELISA assay for GABA, we observed a significant reduction in GABA levels in SCI+KDS 1d/Re, while SCI+V showed aberrant GABA levels at PI 10w (Supplementary Fig. 7j, k). These results indicate that the augmented GABA originates from the increased tonic release of GABA from reactive astrocytes in SCI.

**Fig. 3.**
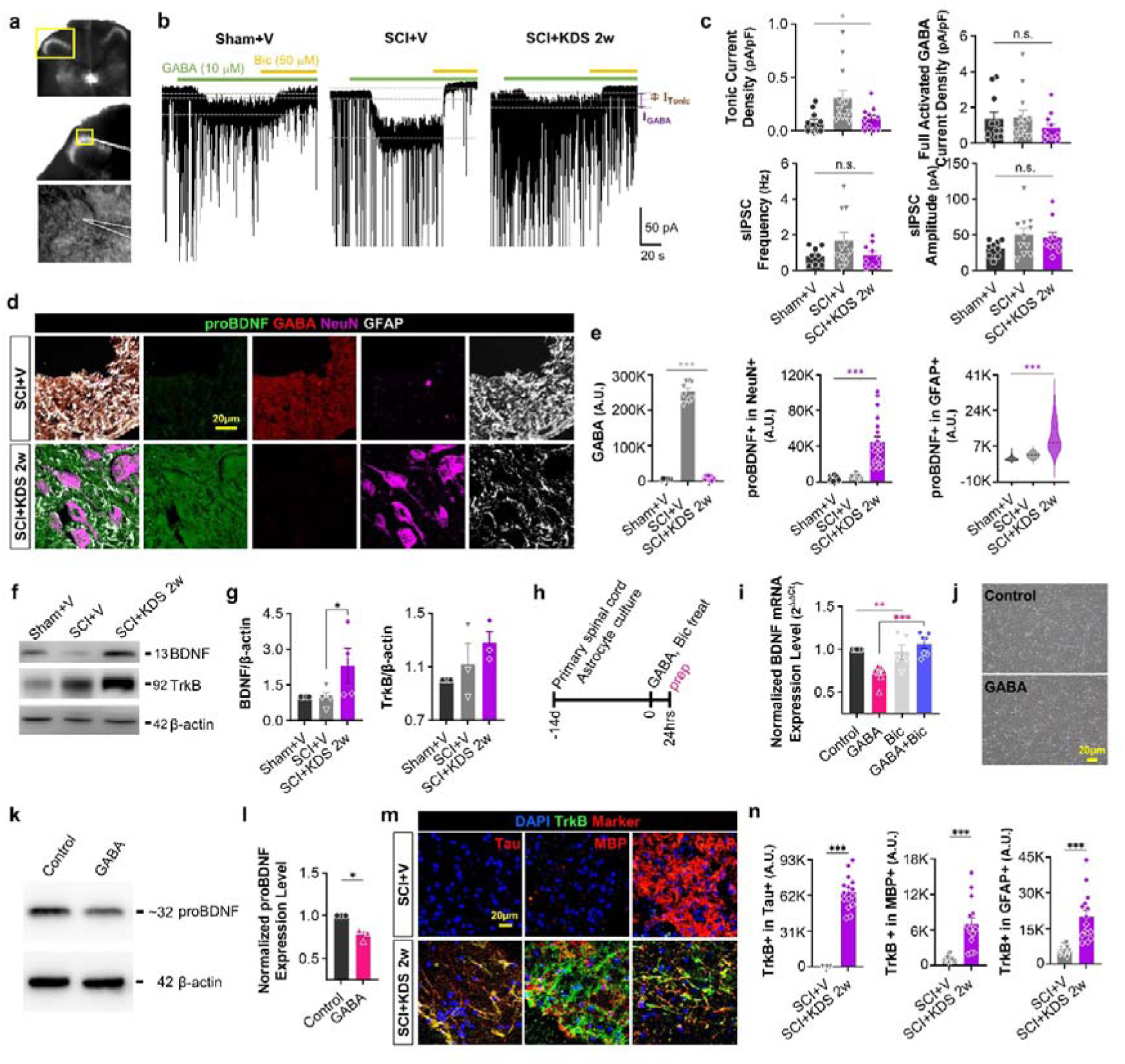
MAOB inhibition with KDS2010 reduces astrocytic GABA level and enhances proBDNF and TrkB expression after SCI. **a** Differential interference contrast (DIC) images of the dorsal horn of the spinal cord (top and middle) and a magnified view of a whole-cell patch-clamped lamina II neuron (bottom). The yellow boxes indicate the magnified region of interest. **b** Representative traces of GABA_A_ receptor-mediated tonic GABA current in each group (Sham+V, SCI+V, and SCI+KDS 2w). Dashed lines (gray) and double-headed arrow (purple and brown) indicate baseline shifts (I_GABA_ and I_Tonic_) with bath application of GABA (10 μM, green bar) and bicuculline (Bic, 50 μM, orange bar). **c** Tonic GABA current density (top, left), GABA-induced full activation current density (top, right), frequency (bottom, left) and amplitude (bottom, right) of spontaneous inhibitory postsynaptic currents (sIPSCs) in each group. **d** Confocal images of the injured areas stained with anti-proBDNF (green), GABA (red), NeuN (magenta), and GFAP (white) antibodies at PI 10w in SCI+V and SCI+KDS 2w. **e** Mean intensity of GABA (left), NeuN-positive proBDNF (middle), GFAP-positive proBDNF (right). **f** Western blotting of BDNF and TrkB in Sham+V, SCI+V, and SCI+KDS 2w at PI 10w. **g** Quantification of BDNF (left) and TrkB (right) expression levels in Western blot. β-actin was used as a control for protein amount. **h** Experimental timeline for quantitative real-time PCR with GABA (100 μM) or Bic (50 μM) treatment in primary cultured spinal cord astrocytes at 14 days *in vitro* (DIV). **i** Relative (comparative Ct) BDNF expression level of each drug treatment condition (GABA, GABA+Bic, and Bic). **j** Representative images of control and GABA-treated spinal cord astrocytes. **k** Western blotting of proBDNF in control and GABA-treated condition. **l** Quantification of proBDNF in Western blot. β-actin was used as a control for protein amount. **m** Confocal images of the injured areas stained with anti-TrkB (green), DAPI (blue), and each Tau, MBP, or GFAP (red) antibodies at PI 10w. **n** Mean intensity of TrkB in the Tau-positive neurons (left), MBP-positive oligodendrocytes (middle), and GFAP-positive astrocytes (right).

We then investigated whether proBDNF expression level could be altered by SCI or MAOB inhibition using immunohistochemistry. Remarkably, we found that the intensity of proBDNF in both astrocytes and neurons significantly increased in SCI+KDS 2w along with a significant decrease in GABA expression (Fig. 3d, e), even at different doses and phases of KDS2010 treatment (Supplementary Fig. 6e, f, Supplementary Fig. 7l, m, Supplementary Fig. 8h, i). This was confirmed by Western blot (Fig. 3f, g). The appearance of proBDNF coincided with the disappearance of astrocytic GABA, reinforcing the idea of the reciprocal relationship between GABA and proBDNF. Next, we treated cultured primary spinal cord astrocytes with GABA for 24 hours and found that GABA significantly down-regulated the mRNA expression level of BDNF, which was fully restored by co-treatment with the GABA_A_antagonist bicuculine (Fig. 3h, i). This GABA-induced proBDNF reduction was also observed in the Western blot (Fig. 3j-l). These results support a novel mechanistic insight that astrocytic GABA acts as an upstream regulator of BDNF signaling.

We then assessed the TrkB expression level in Tau-positive neurons, MBP-positive oligodendrocytes, and GFAP-positive astrocytes. We observed that the intensity of TrkB significantly increased in neurons, oligodendrocytes, and astrocytes following MAOB inhibition from the sub-acute phase (Fig. 3m, n). Similar results were observed for TrkB in oligodendrocytes at different doses and phases of KDS2010 treatment (Supplementary Fig. 6g, h, Supplementary Fig. 8l, m). Given the fact that proBDNF is proteolytically converted to mBDNF by extracellular proteases such as tissue plasminogen activator^36,37^, these results imply that MAOB inhibition causes the removal of astrocytic GABA and a coincidental induction of mBDNF-TrkB signaling pathway, which is indispensable for neuroregeneration and functional/tissue recovery after SCI^34,35^.

Additionally, we observed that KDS2010 treatment reduced the expression of CSPGs, such as neurocan and phosphacan, which are known to inhibit regeneration by creating a hostile extracellular environment (Supplementary Fig. 9d-h). These results further support the idea that MAOB inhibition promotes a more favourable environment for neuroregeneration.

### MAOB inhibition with KDS2010 facilitates tissue recovery in a monkey SCI model

After confirming the efficacy of KDS2010 in promoting recovery from spinal cord injury in small animal models, we performed advanced tests in a non-human primate SCI model, followed by phase 1 clinical trial. This phased approach enabled a thorough evaluation of both safety and therapeutic potential of KDS2010. In the non-human primate study, a severe SCI model was generated in cynomolgus macaque (*Macaca fascicularis*), with KDS2010 administered at doses of 3mpk or 10mpk from the acute phase of SCI (Fig. 4a, b). KDS2010 significantly reduced hematoma and cavity formation at the injury site, indicating a reduction in tissue damage (Fig. 4a). Immunohistochemical analysis revealed that MAP2 intensity was significantly increased in SCI+KDS 10mpk and had a tendency toward increase of MAP2 intensity in the SCI+KDS 3mpk group, compared to SCI+V (Fig. 4c, d). Moreover, astrocytic MAOB was significantly reduced in both KDS treatment groups, while increased astrocytic MAOB was observed in SCI+V (Fig. 4c, d), confirming the effect of KDS2010 on inducing signs of neuroregeneration and reducing the astrocyte reactivity.

**Fig. 4.**
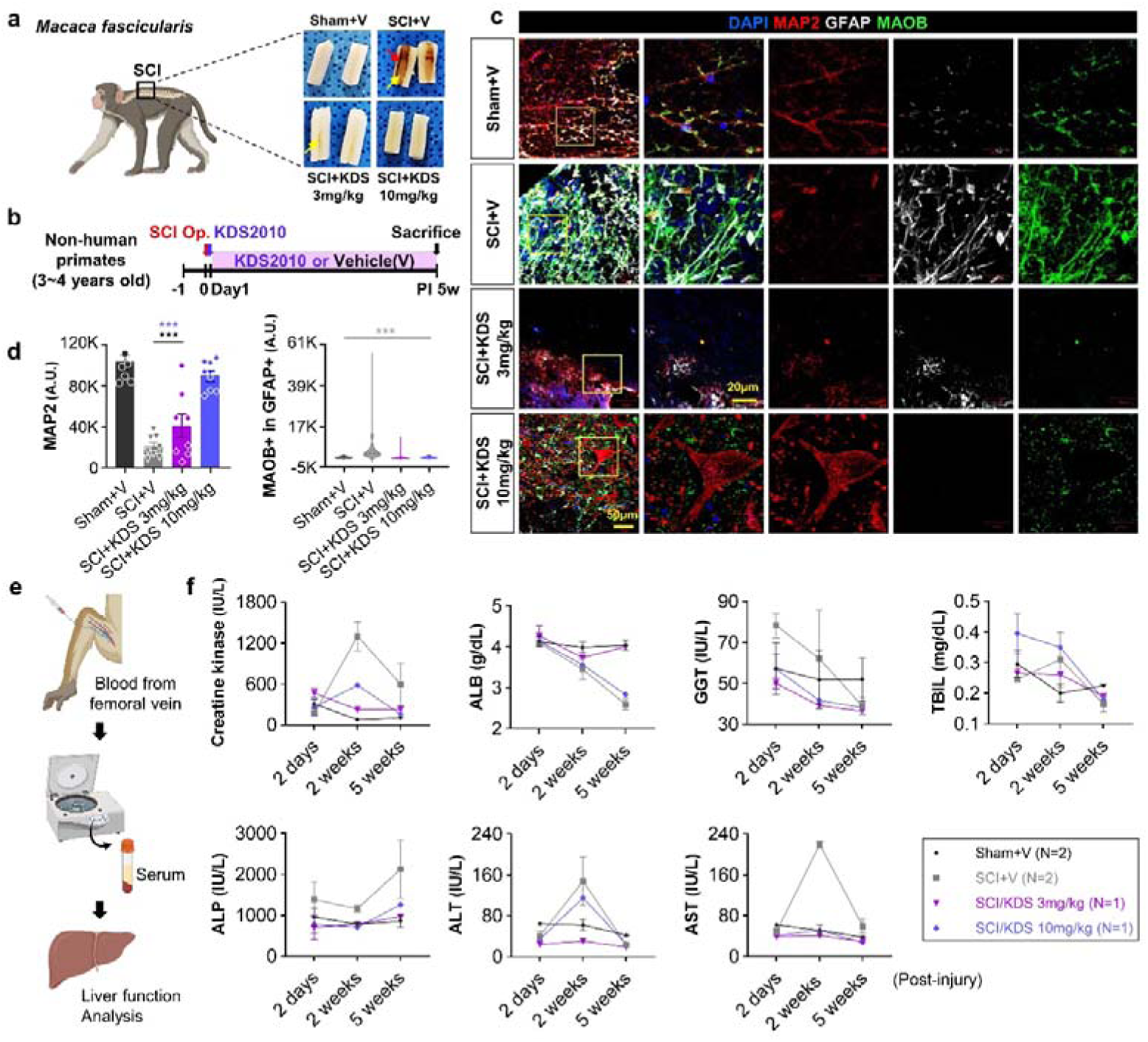
MAOB inhibition with KDS2010 facilitates tissue recovery after SCI in non-human primates. **a** Location of the SCI and representative longitudinal tissue sections at PI 5w in cynomolgus macaque (*Macaca fascicularis*). Red arrows indicate hematoma, and yellow arrows indicate the cavity. **b** Experimental timelines using 3∼4 years old of cynomolgus macaque with the SCI operation and treatment with KDS2010 from 1 day after (acute) SCI. **c** Confocal images of the injured areas stained with anti-MAP2 (red), GFAP (white), and MAOB (green) antibodies, and DAPI (blue) at PI 10w in each group (Sham+V, SCI+V, SCI+KDS 3mpk, and SCI+KDS 10mpk). **d** Mean intensity of the MAP2 (left) and GFAP-positive MAOB (right) in each group. **e** Methodology illustration for blood chemistry analysis with non-human primate SCI model. **f** Blood chemistry and liver function data before and after KDS2010 treatment, showing levels of serum creatine kinase (IU/L), albumin (ALB), gamma-glutamyl transferase (GGT, IU/L), total serum bilirubin (TBIL, mg/dL), alkaline phosphatase (ALP, IU/L), alanine transaminase (ALT, g/dL), and aspartate transaminase (AST, IU/L) at 2 days, 2 weeks, and 5 weeks post-injury.

However, in the blood chemistry and liver function analysis, the 10mpk group exhibited mild signs of hepatic and muscle stress, evidenced by a moderate reduction in albumin (ALB) levels at PI 5w and a transient increase in creatine kinase (CK) and alanine transaminase (ALT) at PI 2w, whereas the 3mpk group showed no signs of stress (Fig. 4e, f). Although CK and ALT levels returned to normal at PI 5w in the 10mpk group (Fig. 4e, f), these findings suggest that prolonged high-dose treatment may require closer monitoring for potential liver toxicity. Taken together, while the 10mpk group showed better recovery than the 3mpk group, 3mpk is considered the No-Observed-Adverse-Effect Level (NOAEL) due to its safer profile.

### KDS2010 exhibits a tolerable safety profile and dose-proportional pharmacokinetics in humans

A phase 1 clinical trial was conducted between September 2022 and May 2024 (Clinical Research Information Service registry no. KCT0008331). A total of 88 subjects were enrolled, with 87 completing the study. One subject withdrew due to a COVID-19 infection during the washout period of the food effect study (Fig. 5a). The 80 young healthy subjects had a mean age of 30.19 years; 77 were men and 3 were women; 41 were Korean and 39 were Caucasian. The 8 elderly male Korean subjects had a mean age of 74.00 years (Supplementary Table 1).

**Fig. 5.**
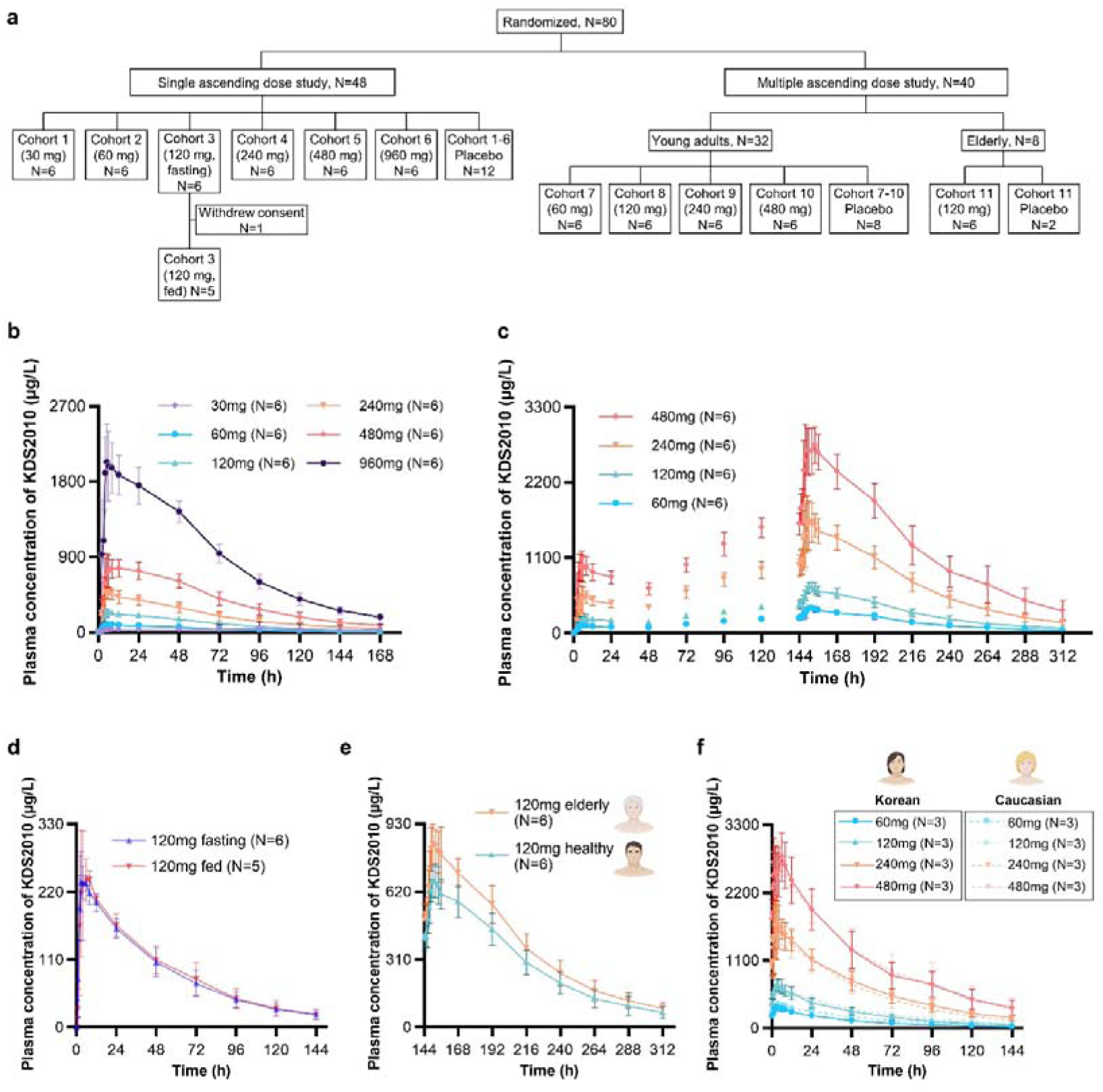
Pharmacokinetic profiles of KDS2010 in human during the phase 1 clinical trial. **a** Subject disposition chart in phase 1 clinical trial. **b, c** Plasma concentration-time profiles of KDS2010 in single ascending dose (**b**) and multiple ascending dose (**c**) study. Dots and error-bars indicate mean concentration and standard deviations, respectively. **d** Plasma concentration-time profiles of KDS2010 following a single 120 mg administration under 10-hour fasted condition and after high-fat meal. Dots and error-bars indicate mean concentration and standard deviations, respectively. **e** Plasma concentration-time profiles of KDS2010 at steady state following 7-day administrations of 120 mg in elderly and young adults. **f** Plasma concentration-time profiles of KDS2010 following administrations of 60, 120, 240, and 480 mg at steady state in Korean and Caucasian subjects.

The safety profile of KDS2010 was tolerable, with no serious adverse events observed. Treatment-emergent adverse events (TEAEs) were reported in 68 cases: 56 TEAEs in 25 subjects (37.9%) on KDS2010 and 12 TEAEs in 6 subjects (27.3%) on placebo (Table 1). The majority of TEAEs were symptoms related to autonomic nervous system including somnolence, hypersomnia, dry eye, conjunctival hyperaemia, and nasal congestion. The incidence of TEAEs tended to increase with dose, and high-fat meal did not affect the frequency of TEAEs. Elderly subjects experienced gastrointestinal symptoms such as constipation, diarrhoea, and abdominal discomfort. The incidence of TEAEs was similar between Korean and Caucasian subjects (data not shown).

**Table 1.**
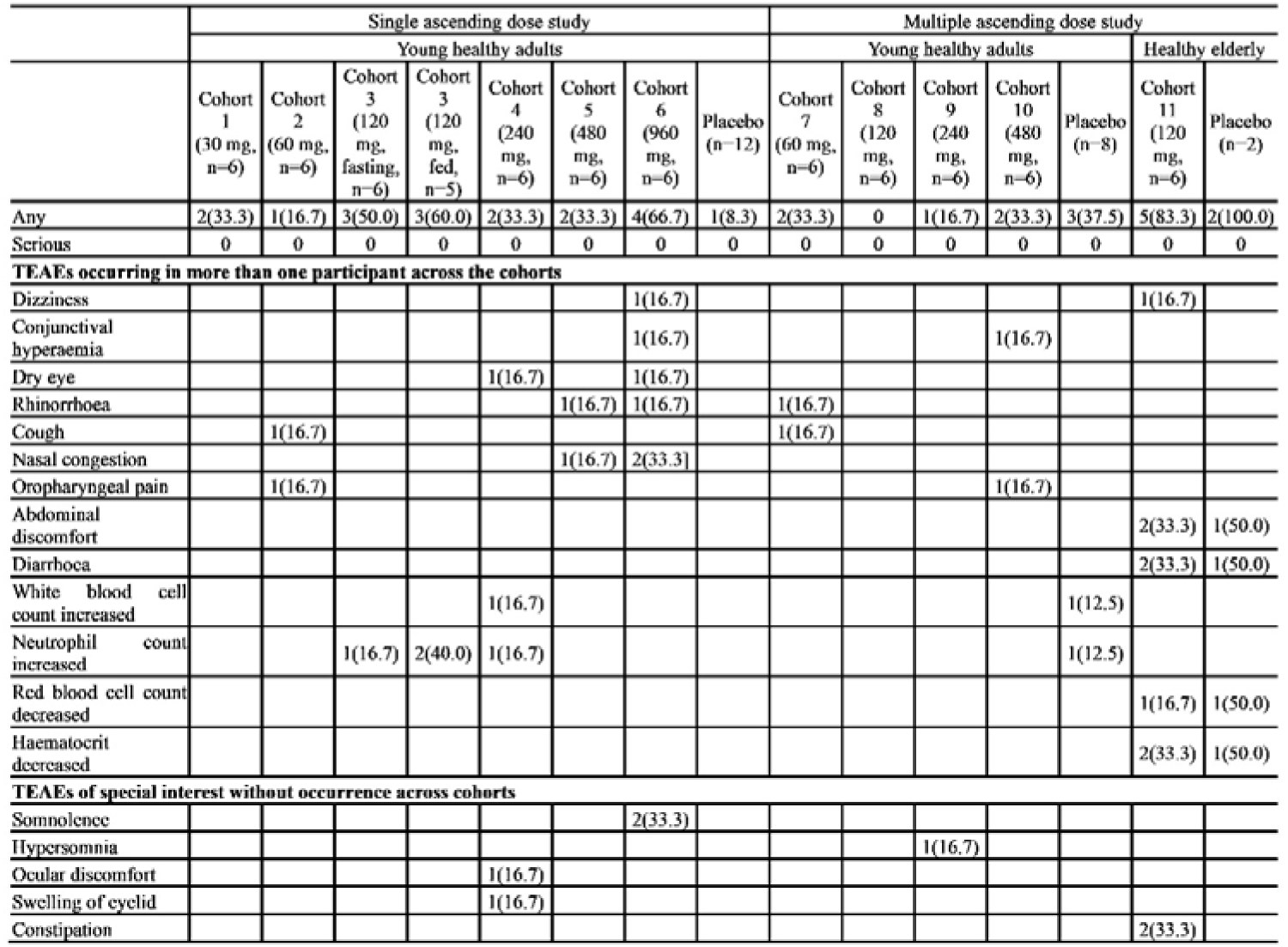
Incidence of treatment-emergent adverse events (TEAEs) in phase 1 clinical trial. All data are presented as the number of subjects (%). The number of randomized participants who received at least one dose of KDS2010 or placebo was indicated as n.

Following a single oral administration, KDS2010 reached its maximum plasma concentration (C_max_) within a median of 3 hours (Fig. 5b, Supplementary Table 2). The concentration declined biexponentially with an average terminal half-life of 44.2 hours. In the 120 mg group, the area under the curve extrapolated to infinity (AUC_inf_) was 13850.7 ng*hr/mL on average. Less than 1% of the unchanged drug was excreted in the urine. The metabolite showed, on average, an 8.54-fold higher systemic exposure than KDS2010. A high-fat meal did not affect the systemic exposure (C_max_ and AUC_inf_) after administration of KDS2010 120 mg (Fig. 5d). After 7 days of multiple oral administrations, the AUC during the dosing interval at steady state (AUC_τ,ss_) was, on average, 3.26-fold higher than that observed on day 1 (Fig. 5c). The systemic exposure (AUC_inf_ and AUC_τ,ss_) increased dose-proportionally from 60 mg to 480 mg. Elderly subjects exhibited approximately 25% higher exposure: geometric mean ratio (GMR) of C_max_ and AUC_τ,ss_ were 1.2521 and 1.2525, respectively (Fig. 5e). However, the inactive metabolite was produced less in the elderly compared to healthy young subjects, with an average metabolic ratios of 4.69 and 7.69 at steady state, respectively. The pharmacokinetic profiles were similar between Korean and Caucasian subjects (Fig. 5f).

## DISCUSSION

In this study, we discovered that GABA from scar-forming reactive astrocytes acts as a brake on the CNS repair system in severe SCI animal models. We could remove this brake through genetic or pharmacological inhibition of MAOB, leading to significant functional and tissue recovery after SCI. Notably, the selective MAOB inhibitor, KDS2010, causes robust neuroregeneration with neuronal proliferation near the injury site, along with a strong induction of astrocytic proBDNF and neuronal TrkB, which can contribute to regenerative process. Importantly, the safety of this drug is confirmed in the phase 1 clinical trial. These novel findings provide a comprehensive understanding of the intricate interplay between astrocytes and the regenerative processes specific to SCI.

BDNF functions as the mature form, mBDNF, which is processed from proBDNF by tissue plasminogen activator (tPA) in the extracellular space or prohormone convertase 1/3 (PC1/3) in intracellular vesicles^38,39^. Upon binding to TrkB, mBDNF activates Akt (Protein kinase B) through PI3K signaling in post-synaptic cells^40,41^, enhancing neuroregeneration by improving cell survival, differentiation, and synaptic plasticity while inhibiting apoptosis^42–44^. Moreover, numerous reports emphasize its importance in SCI^34,45,46^. Thus, it is not surprising that the BDNF-TrkB signaling is critically involved in neuroregeneration and functional recovery after SCI. However, the precise molecular mechanism of how proBDNF expression can be induced in astrocytes is still not clear. So far, we only know that GABA can suppress the BDNF mRNA and proBDNF protein expression in spinal cord astrocytes (Fig. 3h-l). It has been reported that many blood-borne serine proteases rush into the injury sites and activate the protease-activated receptor 1 (PAR1) in astrocytes^47,48^. These raise the possibility that PAR1 activation can induce BDNF mRNA and proBDNF protein expression. Indeed, PAR1 activation has been shown to induce BDNF release in blood platelets^49^. Thus, future experiments are needed to identify the BDNF-inducing factors and determine the precise mechanism of BDNF induction during SCI.

In most cases of MAOB inhibition with KDS2010, it is sufficient to promote recovery after SCI and reduce reactive astrogliosis. However, when KDS2010 was administered from the acute phase, it failed to achieve this in some animals (Supplementary Fig. 7). These results raise intriguing questions of why the MAOB inhibitor works in most animals but not in some animals. We have previously reported that DAO, as an alternative pathway to MAOB, can produce GABA^25,29^ and, MAOB-dependent astrocytic H_2_O_2_ causes reactive astrogliosis in animal models of Alzheimer’s disease^25^. Therefore, it is possible that MAOB might be dominantly expressed and active over DAO in most cases of SCI, whereas in other cases, DAO might be dominant over MAOB and continue to produce H_2_O_2_ to enhance reactive astrogliosis and GABA to inhibit proBDNF even in the presence of KDS2010. Future work is needed to explore these exciting possibilities using effective DAO inhibitors.

The translational relevance of our findings is particularly strong, as the effects of MAOB inhibition are consistently observed across species and various SCI recovery phases. The robust preclinical evidence supporting KDS2010’s efficacy in both rodents and non-human primates suggests that its clinical application could yield substantial benefits for SCI patients. However, some limitations must be acknowledged. While our study establishes a strong correlation between neuroregeneration and functional recovery, future experiments using chemogenetic or optogenetic tools could help clarify the causal relationship between these processes, providing deeper insights into the precise mechanisms by which KDS2010 promotes CNS repair.

Based on the preclinical studies and phase 1 clinical trial, the effective dose of KDS2010 is anticipated to be 120 mg. The pharmacologically active dose (PAD) was 10mpk for 8 weeks for SD rats and 3mpk for 5 weeks for cynomolgus macaques. In each animal model, the systemic exposure (AUC_last_) was approximately 3000 and 13000 ng*hr/mL, respectively. The dose range for phase 1 clinical trial was determined using a pharmacokinetic model based on preclinical plasma concentration data. The model consisted of two compartments with a clearance proportional to the 0.75^th^ power of body weight and volume of distribution to the body weight. In healthy humans, the administration of 120 mg of KDS2010 achieved the target therapeutic exposure of the preclinical PAD, with an AUC_inf_ of 13850.7 ng*hr/mL on average.

At higher doses (240-960 mg), TEAEs related to the autonomic nervous system were observed. MAO oxidatively deaminate endogenous monoamines, including norepinephrine, dopamine, and serotonin^50^. Consequently, neurotransmitter imbalance has been suggested as a potential cause of adverse events associated with MAO inhibitors^51^. Sleep disturbances, dry mouth, and constipation observed in this study have also been reported with other traditional MAOB inhibitors^52–54^. The autonomic nervous system-related adverse events were reported to be reversible^55^, consistent with the findings of this study.

In phase 2 clinical trial, functional recovery effect of KDS2010 120 mg will be explored in SCI patients. When recruiting the SCI patients, stratification by injury severity and phase (acute, sub-acute, and chronic) could be helpful. Standardized diagnostic tools, such as the American Spinal Injury Association (ASIA) Impairment Scale, will help accurately assess the degree of injury and functional impairment. Primary endpoints should focus on functional recovery, including motor and sensory improvements, while secondary endpoints could assess pain, quality of life, and autonomic function. Integrating rehabilitation therapies in the study will be crucial to understanding how KDS2010 works with standard care, potentially maximizing recovery outcomes. By addressing these factors, the phase 2 trial will provide vital insights into the clinical efficacy and broader therapeutic potential of KDS2010 for SCI patients.

In summary, we have traversed the unexplored territory of scar-forming reactive astrocytes and identified MAOB-dependent GABA as the key molecular brake that masks the regenerative process of the mammalian spinal cord after injury. Notably, KDS2010 has shown promise in preclinical studies involving rodent and non-human primate models and was confirmed as safe in the phase 1 clinical trial. We hope that the newly developed concepts and tools related to SCI will be therapeutically helpful in clinical settings where there is currently no available option.

## MATERIALS AND METHODS

### Animals

All animals were housed in facilities accredited by the Association for Assessment and Accreditation of Laboratory Animal Care (AAALAC) at each center. All experimental procedures with mice and rats were conducted following protocols approved by the Institutional Animal Care and Use Committee (IACUC) of Yonsei University (Protocol No. 2016-0027; Seoul, Republic of Korea) and the Institute for Basic Science (Protocol No. IBS-2023-004; Daejeon, Republic of Korea). *B6;129S-Maob^tm1Shih^/J* (MAOB KO; RRID:IMSR_JAX:014133, The Jackson Laboratory, Bar Harbor, ME, USA) mice, which are X-linked *MAOB* gene deficient mouse, were backcrossed with 129S4/SvJaeJ (The Jackson Laboratory) for more than 10 generations and were thought as congenic to 129S4/SvJaeJ. They (8-week-old; 20g ± 3g) were used for SCI experiments. 129S4/SvJaeJ were used as control. B6-*Maob^em1Cjl^*/Ibs (*Maob* floxed) was generated from the Cyagen Biosciences (Guangzhou, China). Female heterozygous *Maob* floxed mice (B6-*Maob^em1Cjl^*/Ibs) were crossed with male transgenic hGFAP-CreER^T2^ (B6-Tg(GFAP-cre/ERT2)13Kdmc; MGI:3712447) mice to generate male *Maob* floxed::hGFAP-CreER^T2^ as previously described^56^. Adult (7-week-old) *Maob* floxed::hGFAP-CreER^T2^ mice were treated with the tamoxifen at 100 mg/kg once per day for 5 days by intraperitoneal (IP) injection to generate astrocyte-specific MAOB cKO mice. Tamoxifen was dissolved in sunflower oil containing 10% ethanol at concentration of 20 mg/ml. For control mice, same amount of sunflower oil was injected to the same age of *Maob* floxed::hGFAP-CreER^T2^ mice. One week after injection, they (8-week-old; 20g ± 3g) were used for SCI experiments. For astrocyte-specific transgene expression of human MAOB in C57BL/6-Tg(Gfap-rtTA,tetO-MAOB,-lacZ)1Jkan/J (GFAP-MAOB; The Jackson Laboratory, 8-week-old; 20g±3g) mice, animals were fed with doxycycline at 3000 ppm provided in pre-mixed Purina chow (Research Diets) for three weeks^57^. 129S4/SvJae-Bdnf*^tm3Jae^*/J (*Bdnf* floxed; The Jackson Laboratory) were backcrossed with C57BL/6J for more than 10 generations and were thought as congenic to C57BL/6J. To generate astrocyte-specific BDNF cKO mice, B6-*Bdnf* floxed mice were crossed with hGFAP-CreER^T2^ or Aldh1l1-Cre/ER^T2^. We used C57BL/6J strain of Aldh1l1-Cre/ER^T2^ by backcrossing B6N.FVB-Tg(Aldh1l1-cre/ERT2)1Khakh/J (The Jackson Laboratory) with C57BL/6J for more than 10 generations. Tamoxifen or saline was injected using the same procedures as in the astrocyte-specific MAOB cKO experiment. C57BL/6J mice (8-week-old; 20g ± 3g ; The Jackson Laboratory) were used for SCI experiments with L655,708. Sprague-Dawley (SD) rats (Orient Bio, Republic of Korea; 8-week-old; 200g ± 20g) were used for SCI experiments. Additionally, female cynomolgus monkeys (3 kg; aged 3–4 years) were used for SCI experiments. All procedures involving monkeys were approved by the IACUC of the Korea Institute of Toxicology (KIT) (Approval No. B216074; Jeongeup-si, Republic of Korea).

### Primary mouse spinal cord astrocyte culture

Primary cultured astrocytes were prepared from spinal cord of C57BL/6J mouse (Jackson Laboratory) pups (P0-P2). Entire spinal cord was dissected with removing all dorsal root ganglia, and meninges covering the outer surface of the spinal cord were removed. Dissected spinal cord was minced and dissociated into single cell suspension by trituration. Cells were plated onto cell culture dish coated with 0.1 mg/ml poly D-lysine (PDL, P6407, Sigma-Aldrich). Cells were cultured in Dulbecco’s modified Eagle’s medium (DMEM, 10-013, Corning) supplemented with 25 glucose, 4 L-glutamine, 1 sodium pyruvate (in mM), 10% heat-inactivated horse serum (26050-088, Gibco), 10% heat-inactivated fetal bovine serum (10082-147, Gibco) and 100 units/ml penicillin–streptomycin (15140-122, Gibco). Cultures were maintained at 37 °C in a humidified 5% CO_2_ incubator. On 3 days in vitro (DIV), cells were vigorously washed with repeated pipetting and the media was replaced to get rid of debris and other floating cell types. During maintaining the culture before use, the media was replaced every 3-4 days.

### Establishment of SCI crush model

For the establishment of mouse and rat SCI models, we followed methods consistent with previous studies^58–60^. Injury was induced at thoracic level 9 in rats for 10 seconds and at thoracic level 10 in mice for 3 seconds. Post-surgical care included administering antibiotics to support recovery and prevent infection, in line with established protocols. Each “Sham” group or “SCI” group indicates animals operated by only laminectomy or SCI, respectively.

For the non-human primate SCI model, female cynomolgus macaque were anesthetized with Zoletil 50 (8–10 mg/kg; Virbac Korea) and isotropy 100. A laminectomy was performed at thoracic level 8 under X-ray guidance, and the SCI model was established using a 60 g vessel clip applied for 10 minutes. After surgery, the monkeys were monitored until full recovery, with precautions taken to prevent hypothermia and dehydration. For pressure ulcers, antibacterial drugs were administered, and sugar therapy was applied as needed.

### Drug delivery

Three types of drugs were used in this study; (S)-2-(((4’-Trifluoromethylbiphenyl-4-yl)methyl)amino)propanamide methanesulfonate (KDS2010) as a reversible MAOB inhibitor^12^, ethyl (S)-11,12,13,13a-tetrahydro-7-methoxy-9-oxo-9H-imidazo[1,5-a]pyrrolo[2,1-c][1,4]benzodiazepine-1-carboxylate (L655,708; L9787, Sigma-Aldrich, St.Louis, MO, USA) as an inverse agonist at the α5 subtype of the benzodiazepine binding site on the GABA_A_ receptor^61^, and 2-hydroxy-5-[2-(4-trifluoromethylphenyl) ethylamino] benzoic acid (AAD2004) as a ROS scavenger. KDS2010 was dissolved in distilled water and administered by drinking *ad libitum* to the SD rats and cynomolgus macaques. KDS2010 was given at a dose of 10 mg/kg from the acute (PI 1 day), sub-acute (PI 2w), and chronic phase (P 6w). To assess dose-dependent effects, additional doses of 20 mg/kg, and 30 mg/kg were administered starting at 2 weeks post-injury. L655,708 was prepared at a concentration of 5mM in DMSO and diluted in 0.9% saline at a dose of 200 µg/kg/day, and administered by i.p. injection to the C57BL/6J mice from PI 2w. AAD2004 was dissolved in dimethyl sulfoxide (DMSO, 0.4%, final concentration 2 mM diluted in phosphate-buffered saline) and administered by i.p. injection at a dose of 3.3 mg/kg/day to the SD rats from PI 2w.

### Viruses and stereotaxic surgery

PCR was employed to amplify the promoter region of the rat *Mki67* gene, encoding the proliferation marker protein Ki67, using rat genomic DNA (69238-3CN, Novagen) as a template. The amplified sequences comprised the 1.4 kb putative rKi67 promoter region, including sequences upstream of the transcription start site. That Ki67 (rKi67) promoter, EGFP, and Cre recombinase genes were incorporated into the AAV-rKi67-EGFP-Cre viral vector, which was then cloned into the AAV-MCS (multiple cloning site) expression vector (VPK-410, Cell Biolabs, Inc.) using the In-Fusion method (639649, Clontech). The AAV-EF1α-DIO-mCherry viral vector was purchased (47636, Addgene). These viral vectors were subsequently purified by iodixanol gradients at the KIST Virus Facility.

For validation of Ki67 promoter, we injected 2 μl of AAV-rKi67-EGFP-Cre (4.58 × 10^13^ genome copies per milliliter (GC/ml)) and AAV-Ef1α-DIO-mCherry (5.56 × 10^13^ GC/ml) with a 1:1 ratio bilaterally into the dentate gyrus (AP = −3.8 mm, ML = 1.8 mm, DV = 3.5 mm) of SD rats (7-week-old). 2 weeks after virus injection, rats were sacrificed for immunohistochemistry.

For labeling the proliferating and proliferated cells after SCI using Ki67 promoter, we injected 3 μl of AAV-rKi67-EGFP-Cre (4.58 × 10^13^ GC/ml) and AAV-Ef1α-DIO-mCherry (5.56 × 10^13^ GC/ml) with a 1:1 ratio into the spinal cord at two locations (1 mm proximal and distal to the injured site) of rat SCI model 2 weeks after SCI. At PI 10w (8 weeks after virus injection), rats were sacrificed for immunohistochemistry.

### Behavioral test

Behavioral test was performed every week for total of 11 weeks (1 week before and 10 weeks after the surgical operation) to assess locomotor recovery. The Basso, Beattie, and Bresnahan (BBB) motor score^27^ or Basso Mouse Scale (BMS) score^17^ was used to evaluate the quality of hind limb movement during open field locomotion in rats or mice, respectively. In the first recovery phase, the range of joint movement and the presence of the foot closure on the floor were checked. In the second phase, recovery of weighted stepping was checked. In the third phase, gait coordination and tail movement were assessed. These scores are “ordinal and the magnitude of behavioral change between ranks may not be consistent,” as originally described^17^. To perform the group allocation in a blinded manner during data collection, animal preparation and behavior test were performed by at least two different investigators.

In experiments involving acute KDS2010 administration and transgenic mouse models, animals were grouped based on similar average BBB or BMS scores one day post-injury. In the BBB locomotor test with drug administration from PI 2w, animals were initially subjected to injury, then divided into two groups at PI 2w based on similar averaged BBB scores. Subsequently, these groups were treated with either the vehicle (SCI+V) or the drug (SCI+drug) from PI 2w. To exclude incompletely generated severe rat SCI animals, we established criteria: a score difference of over 4 between PI 1w and 2w, and a score exceeding 8 at PI 2w. In the BMS locomotor test of L655,708 experiment, we excluded animals that did not score 0 one day after SCI, as previously described in a report^62^.

The ladder rung test was performed with an apparatus featuring a 100 cm horizontal ladder with 1cm spacing between rungs. The functional recovery was measured by the percentage of hindlimb steps without slipping as rats walked along a horizontal ladder, as previously reported^28,63^.

### Immunohistochemistry

Immunohistochemistry was performed as previously described^64^. Tissue sections were blocked and treated with primary antibodies. After overnight incubation at 4°C, species-specific fluorescent secondary antibodies were applied for visualization. Tissue sections were then stained with DAPI during mounting. This protocol allowed detailed assessment of various cellular markers involved in SCI and recovery processes.

The primary antibodies included chicken anti-MAP2 (1:2000, ab5392, Abcam), goat anti-GFAP (1:1000, ab53554, Abcam), rabbit anti-MAOB (1:200, ab175136, Abcam), mouse anti-GABA (1:1000, ab86186, Abcam), chicken anti-GFP (1:500, ab13970, Abcam), rabbit anti-mCherry (1:500, ab167453, Abcam), rabbit anti-Ki67 (1:500, ab16667, Abcam), rabbit anti-NeuN (1:800, ab177487, Abcam), chicken anti-proBDNF (1:200, AB9042, Millipore), rabbit anti-TrkB (1:100, ab18987, Abcam), chicken anti-Tau (1:200, ab75714, Abcam), chicken anti-Myelin basic protein (1:400, ab134018, Abcam), rabbit anti-beta III tubulin (Tuj1) (1:2000, ab18207, Abcam), and rabbit anti-DAO (1:500, ARP41908_P050, Aviva Systems Biology).

The secondary antibodies included FITC-donkey anti-mouse IgG (H+L) (1:150, 715-095-151), Cy^TM^3-donkey anti-mouse IgG (H+L) (1:600, 715-165-151), FITC-donkey anti-rabbit IgG (H+L) (1:150, 711-095-152), Alexa Fluor 594-donkey anti-rabbit IgG (H+L) (1:500, 711-585-152), Alexa Fluor 647-donkey anti-rabbit IgG (H+L) (1:500, 711-605-152), Cy^TM^3-donkey anti-chicken IgG (H+L) (1:600, 703-165-155), DyLightTM405-donkey anti-chicken IgY++ (H+L) (1:600, 703-476-155), Alexa Fluor 488-donkey anti-chicken IgY (IgG) (H+L) (1:500, 703-545-155), Alexa Fluor 594-donkey anti-chicken IgY (IgG) (H+L) (1:500, 703-585-155), Alexa Fluor 488-donkey anti-goat IgG (H+L) (1:500, 705-545-003), and Alexa Fluor 647-donkey anti-goat IgG (H+L) (1:300, 705-606-147). All secondary antibodies were purchased from Jackson ImmunoResearch.

The fluorescence images were observed under a confocal laser scanning microscopy (LSM700 or LSM710, Carl Zeiss), and analyzed using the ImageJ (NIH) software. For visualizing the mCh-positive signals of proliferating or proliferated cells, Z-stack images acquired in 1-μm steps were processed for three-dimensional (3D) surface rendering of mCh-positive signals using IMARIS software (version 9.0.1, Oxford Instruments).

### Myelin analysis

Myelin integrity and structure were assessed using Eriochrome Cyanine (EC)^65^, Luxol Fast Blue (LFB)^64^, and Toluidine Blue (TB) staining, along with Transmission Electron Microscopy (TEM)^64–66^. All procedures were performed following previously established protocols.

### Tonic GABA current recording

Prior to recording, spinal cords were harvested from rats that were anesthetized with a mixture of Zoletil and Xylazine. The dura mater of the isolated spinal cords was eliminated in the ice-cold recovery solution containing N-Methyl-D-glucamine (NMDG) (NMDG-recovery solution): 93 NMDG, 2.5 KCl, 1.2 NaH_2_PO_4_, 30 NaHCO_3_, 20 HEPES, 25 D-(+)-Glucose, 5 sodium ascorbate, 2 thiourea, 3 sodium pyruvate, 10 MgSO_4_, 0.5 CaCl_2_ (in mM), pH 7.4. The slices (400 μm in thickness were first incubated at 32°C for 15 min in NMDG-recovery solution, then incubated at 22±2°C for at least 1 h in normal artificial cerebrospinal fluid (aCSF) solution: 130 NaCl, 24 NaHCO_3_, 1.25 NaH_2_PO_4_, 3.5 KCl, 1.5 CaCl_2_, 1.5 MgCl_2_, and 10 D-(+)-glucose (in mM), pH 7.4. For the recordings, the slices were transferred to a recording chamber that was continuously perfused with aCSF solution. Whole-cell patch-clamp recordings were performed on the neurons of lamina II in the dorsal horn of the spinal cord within 2 cm of the injury. The holding voltage was -60 mV. The pipette resistance was typically 6–8 MΩ and the pipette was filled with an internal solution: 135 CsCl, 4 NaCl, 0.5 CaCl_2_, 10 HEPES, 5 EGTA, 4 Mg-ATP, 0.3 Na_2_-GTP, and 10 QX-314 (mM), pH adjusted to

7.2 with CsOH (278–285 mOsmol/kg). Electrical signals were digitized and sampled at 50-ms intervals with a Digidata 1440A and Multiclamp 700B amplifier (Molecular Devices) using pCLAMP 10.4 software (Molecular Devices). Data were filtered at 2 kHz. D-AP5 (50 μM; 0106, Tocris), CNQX (20 μM; 0190, Tocris), GABA (10 μM; A2129, Sigma-Aldrich), and (-)-bicuculline methobromide (50 μM; BIC; 0109, Tocris) were treated for measuring tonic GABA current and fully-activated GABA current as previously described^11^. The measurement was performed using the Clampfit 10.4 program (Molecular Devices). Every current was divided by the capacitance of each cell to calculate the current density. The frequency and amplitude of spontaneous inhibitory post-synaptic currents (sIPSCs) before GABA application were measured by Mini Analysis Program (Synaptosoft). For the analysis, data points were excluded when the membrane capacitance of the cell was under 5 pF. Finally, Grubb’s test was performed to identify statistical outliers.

### Quantitative real-time PCR

Primary mouse spinal cord astrocytes were treated with GABA (100 μM, A2129, Sigma-Aldrich) and/or bicuculline (50 μM; 0109, Tocris). Approximately 24 hours after drug treatment, total RNA was extracted by using RNA isolation kit (314-150, GeneAll), and cDNA was synthesized by using reverse transcriptase (18080-044, Invitrogen). SYBR-green (4309155, Applied Biosystems) was used to perform quantitative real-time PCR. The following sequences of primers^67^ were used for quantitative real-time PCR. Mouse BDNF forward: 5’-AAAGTCCCGGTATCCAAAGGCCAA-3’; Mouse BDNF reverse: 5’-TAGTTCGGCATTGCGAGTTCCAGT-3’. Mouse GAPDH forward: 5’-TGATGACATCAAGAAGGTGGTGAAG-3’; Mouse GAPDH reverse: 5’-TCCTTGGAGGCCATGTAGGCCAT-3’.

### Western blot and ELISA

Approximately 24 hours after drug treatment, primary mouse spinal cord astrocytes or spinal cord tissues were lysed in RIPA buffer (MB030-0050, ROCKLAND) containing protease and phosphatase inhibitors (1861280, Thermo Fisher Scientific). Protein concentration was measured using the Pierce™ BCA Protein Assay Kit (Thermo Fisher Scientific). Each sample was then used separately for Western blot or GABA ELISA analysis.

Western blot analysis was performed following established protocols^68,69^, with 20 μg of spinal cord tissue or 10 μg of astrocyte protein samples loaded onto SDS-PAGE gels, transferred to PVDF membranes, and blocked with 5% non-fat milk. Membranes were incubated overnight at 4°C with primary antibodies, including rabbit anti-MAOB (1:1000, Abcam), rabbit anti-BDNF (1:1000, Abcam), mouse anti-TrkB (1:1000, R&D Systems), and rabbit anti-β-actin (1:1000, Cell Signaling Technology). Detection was achieved using species-specific secondary antibodies conjugated with horseradish peroxidase (HRP), including anti-rabbit IgG, HRP-linked (1:2500 or 1:5000, 7074, Cell Signaling Technology), anti-mouse IgG, HRP-linked (1:2500, 7076, Cell Signaling Technology), and rabbit anti-goat IgG, HRP-linked (1:2500, ab6741, Abcam). Bands were visualized using enhanced chemiluminescence, and band intensity was quantified using ImageJ software.

For ELISA, GABA quantification was performed using an ELISA kit (BA-E-2500, ImmuSmol) according to the manufacturer’s instructions, utilizing 20 μg of protein extract from spinal cord tissues.

### Evaluation of liver function test parameters in non-human primates

Blood samples were collected from the animals via the radial or femoral vein at three time points: 2 days, 2 weeks, and 5 weeks post-injury. All animals were fasted for approximately 16 hours prior to each collection, with water provided *ad libitum*. Approximately 2.0 mL of blood was drawn and placed into tubes without anticoagulant and left at room temperature for at least 90 minutes. The samples were then centrifuged at 3000 rpm for 10 minutes at room temperature to separate the serum. Finally, serum samples were analyzed for creatinine (CREA), alkaline phosphatase (ALP), gamma-glutamyl transpeptidase (GGT), total bilirubin (TBIL), albumin (ALB), alanine aminotransferase (ALT), and aspartate aminotransferase (AST). These biochemical parameters were measured using a TBA 120FR chemistry analyzer (Toshiba Co.).

### Phase 1 clinical trial study design and subjects

A randomized, double-blind, placebo-controlled study of oral KDS2010 was conducted in healthy Korean and Caucasian subjects (Clinical Research Information Service registry no. KCT0008331). All subjects provided consent for this phase 1 clinical trial study. The study comprised 11 cohorts: SAD and MAD studies. SAD cohorts (1-9) received 30-960 mg of KDS2010. Cohort 3 employed a two-period crossover design with a 14-day washout. Participants received a 120 mg dose of KDS2010 after a 10-hour fast in the first period, and following a high-fat meal (over 900 kcal, 35% fat) in the second. This allowed for a direct comparison of KDS2010 pharmacokinetics in fasting versus fed states. MAD cohorts (7-10) received 60-480 mg of KDS2010 daily for 7 days. Cohort 11 comprised elderly subjects who received 120 mg of KDS2010 daily for 7 days.

Subjects were randomized 3:1 to receive KDS2010 or placebo. Eligibility criteria included healthy adults (aged 19-45) for the SAD and MAD cohorts, and healthy elderly subjects (aged 65-85) for cohort 11. The participants were excluded if they met the following criteria: serum aspartate aminotransferase (AST) or alanine aminotransferase (ALT) greater than 60 IU/L, total bilirubin greater than 1.8 mg/dL, creatinine phosphokinase (CPK) levels greater than 405 IU/L, creatinine clearance calculated by the Chronic Kidney Disease Epidemiology Collaboration (CKD-EPI) equation less than 60 mL/min/1.73m². At the time of randomization, all subjects were required to be free from any clinically significant medical conditions. The study was approved by the Seoul National University Hospital institutional review board and conducted in accordance with the Declaration of Helsinki and Good Clinical Practice guidelines.

Subjects who met the eligibility criteria were admitted to the Seoul National University Hospital Clinical Trial Center one day prior to the first dose administration. On the scheduled dosing day, participants received either KDS2010 or placebo, randomly assigned, with 150 mL of water. After administration of the last dose, subjects were discharged 72 hours later and visited the outpatient clinic for follow-up on three consecutive days. Pharmacokinetics and safety were monitored until the end of the study, lasting up to 18 days after the last dose.

The primary endpoint was the incidence of TEAEs by dose group. Any abnormalities in vital signs, electrocardiograms, clinical laboratory tests, or neurological examinations that occurred after KDS2010 administration were recorded as TEAEs. Those deemed to have a causal relationship with the drug were classified as ADRs. Dose escalation proceeded under blinded conditions, contingent on the absence of dose-limiting toxicity, as reviewed by a safety review committee.

Secondary endpoints included pharmacokinetic parameters such as maximum observed concentration (C_max_), time to Cmax (T_max_), area under the concentration-time curve from 0 to infinity (AUC_inf_), AUC during dosing intervals at steady state (AUC_τ,ss_), terminal half-life (t_1/2_), apparent clearance (CL/F), fraction excreted of the unchanged drug (fe), and renal clearance (CL_R_). The parameters related to plasma concentrations were calculated via non-compartmental analysis using the linear-up log-down method in Phoenix WinNonlin software version 8.3 (Certara, US). The parameters related to urine concentrations were calculated using the linear trapezoidal linear interpolation method.

Serial blood and urine samples were collected at the scheduled times and the concentration of KDS2010 and its metabolite was measured. In the SAD study, plasma samples were collected using sodium K2-EDTA tubes at pre-dose, 0.33, 0.75, 1, 2, 3, 4, 6, 8, 12, 24, 48, 72, 96, 120, and 144 hours after dosing. Urine samples were collected up to 72 hours post-dose. In the MAD study, plasma samples were collected at pre-dose, 0.33, 0.75, 1, 2, 3, 4, 6, 8, 12 and 24 hours after the administration of the first dose, and at pre-dose of the third to seventh doses. After the administration of the seventh dose, plasma samples were collected at the same time points as after the first dose, with additional sampling points at 48, 72, 96, 120, and 144 hours post-dose. Urine samples were collected for 24 hours after the administration of the first dose and for 72 hours after the seventh doses.

The concentrations of KDS2010 in plasma and urine were determined using validated liquid chromatography–tandem mass spectrometry (SCIEX API 4000 system). Standard curves for KDS2010 showed linearity within a range of 5-5000 μg/L in plasma samples and 5-2500 μg/L in urine samples. The lower limit of quantification (LLOQ) for KDS2010 was 5 μg/L in both plasma and urine. For the metabolite, standard curves demonstrated linearity within a range of 1-1000 μg/L in both plasma and urine samples, with an LLOQ of 1 μg/L for both.

### Statistical analysis

For all preclinical experiments, data are presented as mean ± standard error of the mean (SEM). For all experiments, data normality was analyzed using a D’Agostino–Pearson omnibus normality test. For data not following normal distribution, a Mann-Whitney test (Two-tailed) or Kruskal–Wallis test with Dunn’s multiple comparison test was performed. Differences between groups were evaluated by one-way analysis of variance (ANOVA) with Tukey’s multiple comparisons tests. For behavioral test, two-way RM ANOVA with Tukey’s multiple comparisons test and linear mixed model were performed for statistical analysis. Grubb’s test was performed to identify statistical outliers in tonic GABA recording experiment. GraphPad Prism 10.3.1 for Windows (GraphPad Software) and SAS 9.4 (SAS Institute Inc., USA) were used for these analyses.

For the clinical trial, dose proportionality of the PK parameters (C_max_, C_max,ss_, AUC_last_, and AUC_τ_) was assessed using power model using the following equation: log (PK parameter) = β0 +β1 log (Dose). The regression slope (β1) and the intercept (β0) were estimated with 95 % confidence interval (CI) using PROC REG in SAS version 9.4. The PK parameters were considered dose-proportional if the 95% CI of the β1 included 1. The effect of a high-fat meal on the PK parameters (C_max_, C_max,ss_, AUC_last_, and AUC_τ_) was evaluated by calculating the geometric mean ratio (GMR) and its 90 % CI. Using a linear mixed effect model, the logarithmized PK parameters were analyzed with the high-gat meal as a fixed effect and inter-individual differences as a random effect. The GMR and its 90 % CI of the PK parameters of the high-fat meal were calculated relative to those in the fasting state. The effect of a high-fat meal was considered negligible if the 90% CI fell within the range of 0.8 to 1.25. In elderly subjects, the same analyses were performed and compared to those of healthy younger adults.

## DATA AVAILABILITY

The data that support the findings of this study are available from the corresponding author upon reasonable request.

## ACKNOWLEGEMENTS

This research was supported by a faculty research grant of Yonsei University College of Medicine (6-2018-0161), the National Research Foundation of Korea (NRF) grant funded by the Korea government (MSIT) (RS-2024-00341308), Center for Cognition and Sociality from Institute for Basic Science (IBS) (IBS-R001-D2), and NeuroBiogen Co., LTD. We thank the clinical trial participants, as well as the clinical trial investigators of SNU.

## AUTHOR CONTRIBUTIONS

H.Y.L., C.J.L., and Y.H. designed the study. H.Y.L., H.-L.L., J.M.L., and J.P. analyzed the data. H.Y.L., H.-L.L., J.M.L., C.J.L., and J.P. wrote the manuscript. H.Y.L. and H.-L.L. performed animal surgeries, behavioral tests, immunohistochemistry, and analyses. J.M.L., S.Y.C. and S.Y.H. conducted additional behavioral tests and immunohistochemistry. G.Y.H., H.K., and J.W. performed *in vitro* analyses. J.M.L. and M.-H.N. performed slice electrophysiology. S.E.L. constructed the Ki67 promoter. E.K.P., B.K.J., E.H.L., S.W.K., and K.D.P. synthesized and analyzed the compounds. H.Y.L. and K.N.K. conducted primate studies. J.P. and S.L. performed the phase 1 clinical trial. M.G.P. created the schematic figure. All authors contributed to discussions of the results.

