## Supplementary Information for "Astrocytic MAOB-GABA axis as a molecular brake on repair following spinal cord injury"

Yoon Ha, MD, Ph.D.

Department of Neurosurgery, College of Medicine, Yonsei University,

Yonsei-ro 50, Seoul, Republic of Korea

C. Justin Lee, Ph.D.

Director of Center for Cognition and Sociality, Institute for Basic Science (IBS),

Expo-ro 55, Daejeon, Republic of Korea

**This file includes:**

**Supplementary Figures 1 to 10**

**Supplementary Tables 1, 2**

**Supplementary Movie 1, 2**

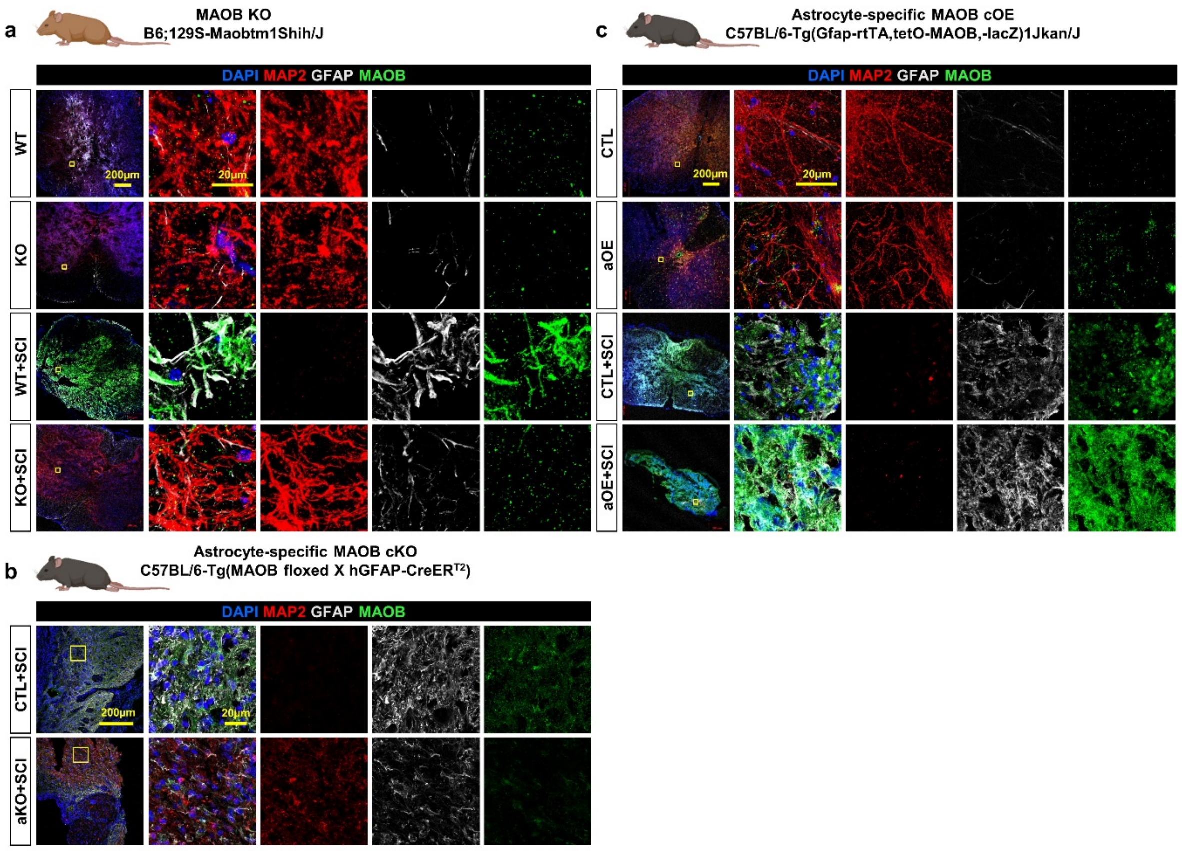


**Supplementary Fig. 1 Immunohistological analysis of astrocyte reactivity and neuronal quantity in MAOB KO, aKO, and aKO mice.**

**a-c** Confocal images of injured area showing individual channels for MAP2 (red), GFAP (white), and MAOB (green) at PI 10w in each group for MAOB KO (**a**), aOE (**b**), and aKO (**c**). Each yellow box in merged images indicates the magnified region of interest.

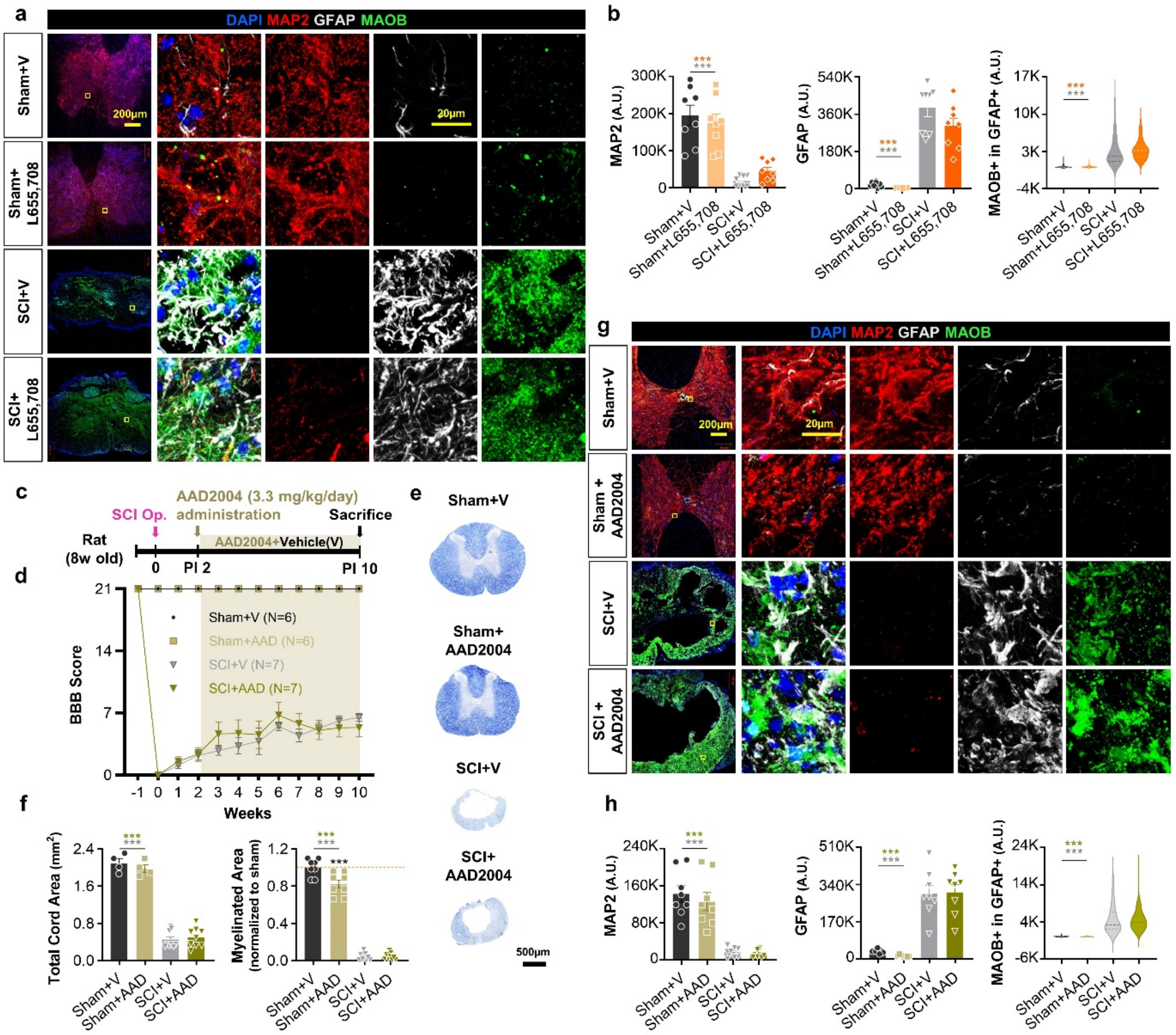


**Supplementary Fig. 2 Inhibiting the α5-containing $GABA_A$ receptor from the sub-acute phase induces recovery after SCI, while scavenging ROS does not.**

**a** Confocal images of the injured area in each group (Sham+V, Sham+L655,708, SCI+V, and SCI+L655,708) stained with anti-MAP2 (red), GFAP (white), and MAOB (green) antibodies at PI 10w. **b** (Left) Compared to SCI+V, SCI+L655,708 showed a tendency toward increased MAP2 intensity. The intensity of GFAP (middle) and GFAP-positive MAOB (right) in SCI+L655,708 had no significant change compared to SCI+V. **c** Experimental timeline using 8-week-old rats with the SCI operation and treatment with the ROS scavenger, AAD2004. **d** BBB score of each group (Sham+V, Sham+AAD2004, SCI+V, and SCI+AAD2004). SCI+AAD2004 did not show any improvement in functional recovery compared to SCI+V. Green shade indicates the duration of AAD2004 administration. **e** EC staining of spinal cord tissues in each group at PI 10w. **f** SCI+AAD2004 did not show any signs of tissue recovery compared to SCI+V. **g** Confocal images of the injured area in each group stained with anti-MAP2 (red), GFAP (white), MAOB (green) antibodies, and DAPI (blue) at PI 10w. **h** The intensity of MAP2 (left) and astrocytic GFAP (middle) and MAOB (right) in SCI+AAD2004 showed no difference compared to SCI+V. Each yellow box in merged images in **a** and **g** indicates the magnified region of interest.

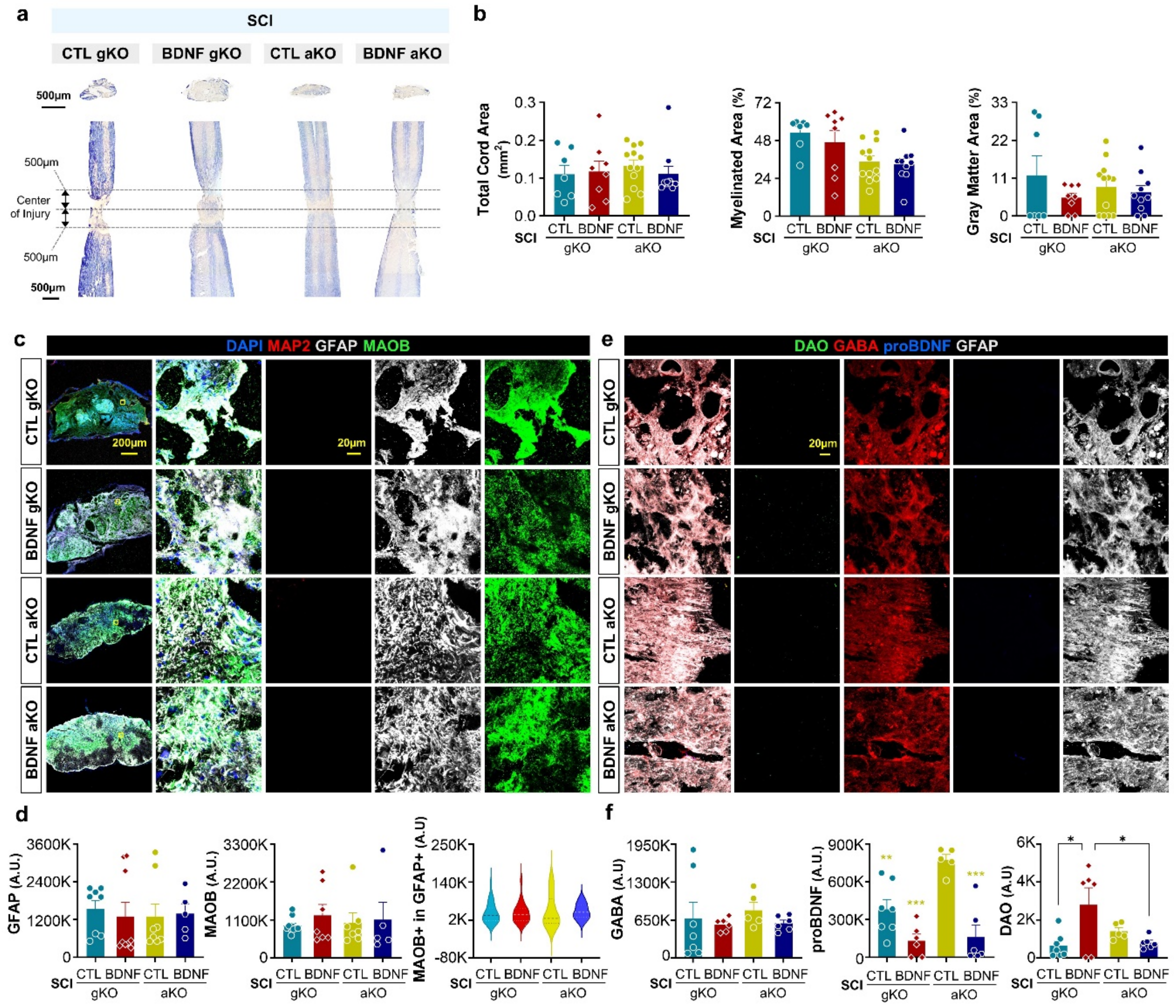


**Supplementary Fig. 3 BDNF signaling is downstream of astrocyte reactivity and astrocytic GABA expression.**

**a** EC staining of cross (top) and longitudinal (bottom) sections of spinal cord tissues in each group (CTL gKO+SCI, BDNF gKO+SCI, CTL aKO+SCI, and BDNF aKO+SCI) at PI 8w. **b** Total cord, myelinated, and grey matter areas did not show any difference among the groups. **c** Confocal images of the injured area in each group stained with anti-MAP2 (red), GFAP (white), MAOB (green) antibodies, and DAPI (blue) at PI 8w. Yellow box indicates the magnified region of interest. **d** The intensity of GFAP (left), MAOB (middle), and GFAP-positive MAOB (right) had no significant change among the groups. **e** Confocal images of the injured area in each group stained with anti-DAO (green), GABA (red), proBDNF (blue), and GFAP (white) antibodies at PI 8w. **f** (Left) GABA intensity showed no significant change among the groups. (Middle) Compared their respective CTL, both BDNF gKO and aKO showed a significant decrease in proBDNF intensity. (Bottom) BDNF gKO showed a significant increase in DAO intensity compared to CTL.

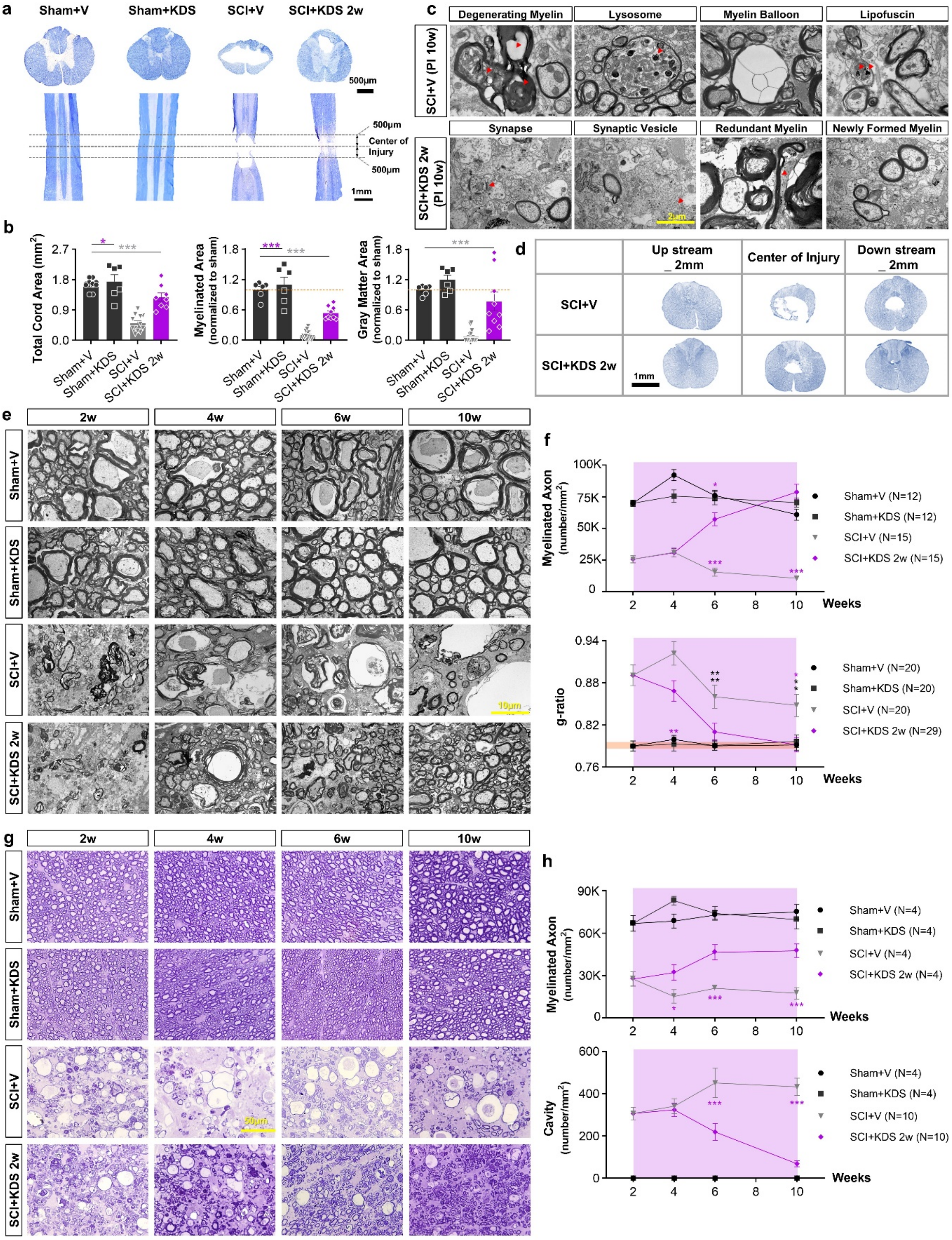


**Supplementary Fig. 4 Signs of remyelination through MAOB inhibition.**

**a** EC staining of cross (top) and longitudinal (bottom) sections of spinal cord tissues in each group (Sham+V, Sham+KDS, SCI+V, and SCI+KDS 2w) at PI 10w. **b** Compared to Sham+V and Sham+KDS,

SCI+V showed significantly reduced total spinal cord, myelinated, and grey matter, along with an enlarged cavity size. All were significantly recovered in SCI+KDS 2w. **c** (Top) Representative TEM images showed evidence of myelin degeneration after SCI (SCI+V), such as degenerating myelin, lysosome, myelin balloon, and lipofuscin, at PI 10w. (Bottom) Representative TEM images showed evidence of neuroregeneration, such as synapse, synaptic vesicle, redundant myelin, and newly formed myelin, at PI 10w, when MAOB was inhibited after SCI (SCI+KDS 2w). **d** Luxol Fast Blue (LFB) staining showed recovery of gray-white matter boundaries and reduced cavity formation in the SCI+KDS 2w compared to SCI+V. **e** TEM images of spinal cord tissues in each group at PI 2, 4, 6, and 10w. **f** Compared to Sham+V and Sham+KDS, SCI+V showed severe loss of myelination (top) and increase of g-ratio (bottom). In contrast, SCI+KDS 2w showed a gradual increase in the number of small-sized myelinated axons (top) and a restoration of g-ratio to the optimal rage of $0.790 \pm 0.005$ (orange shade) at PI 10w. **g** TB staining of spinal cord tissues in each group at PI 2, 4, 6, and 10 week. **h** Compared to Sham+V and Sham+KDS, SCI+V showed severe loss of myelination (top) and the emergence of large cavities (bottom). In contrast, SCI+KDS 2w showed a gradual increase in the number of small-sized myelinated axons (top) and gradual decrease in the density of cavities (bottom). Purple shades in **f** and **h** indicate the duration of KDS2010 administration.

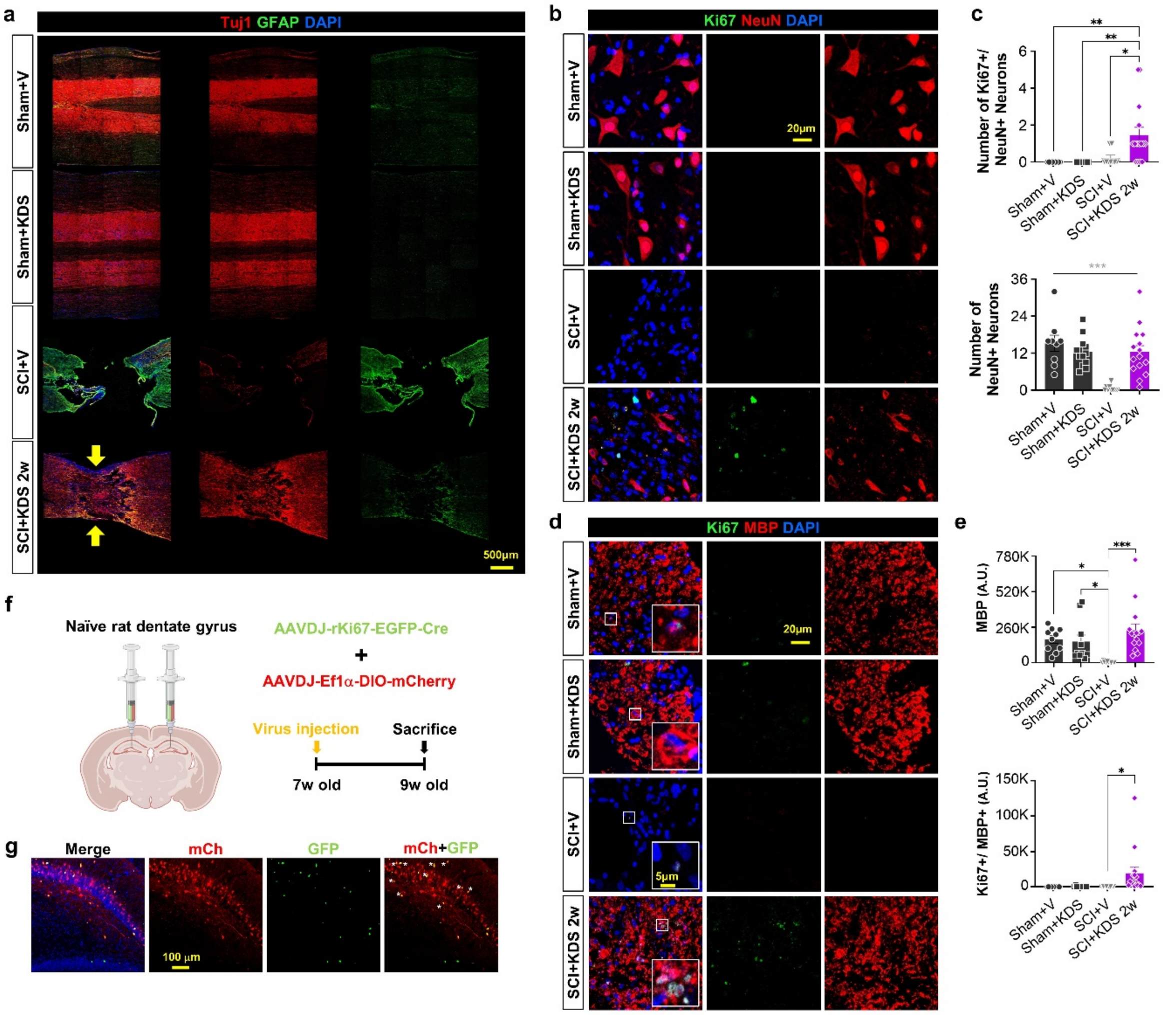


**Supplementary Fig. 5 Increased proliferation of neurons and oligodendrocytes through MAOB inhibition, and validation of the Ki67 promoter in the rat dentate gyrus.**

**a** Confocal images of longitudinal sections of spinal cord tissues stained with anti-Tuj1 (red) and GFAP (green) antibodies at PI 10w in each group with sub-acute phase KDS2010 treatment (PI 2w). Yellow arrow indicates a cavity and atrophy in the center of the injured area. **b** Confocal images of the injured area in each group stained with anti-Ki67 (green), NeuN (red) antibodies, and DAPI (blue) at PI 10w. **c** (Top) Ki67-positive proliferating neurons were significantly increased in SCI+KDS 2w compared to Sham-operated groups and SCI+V. (Bottom) The number of NeuN-positive neurons in SCI+V was significantly reduced in SCI+V, while SCI+KDS 2w showed a significant recovery. **d** Confocal images of the injured area in each group stained with anti-Ki67 (green), MBP (red) antibodies, and DAPI (blue) at PI 10w. **e** (Top) The MBP intensity in SCI+V was significantly reduced compared to Sham-operated groups, while SCI+KDS 2w showed a significant recovery. (Bottom) Ki67-positive proliferating oligodendrocytes were significantly increased in SCI+KDS 2w compared to Sham-operated groups and SCI+V. **f** (Left) Schematics of virus injection strategy targeting the dentate gyrus of rat. (Right) Experimental timeline using 7-week-old rats involving the injection of a mixture of two viruses. **g** Confocal images of the dentate gyrus showing mCh and/or GFP-positive cells stained with anti-GFP (green) antibody and DAPI (blue) at 2 weeks after virus injection. The percentage of proliferating and proliferated cells was approximately 5.14±0.53% (n=3) for 2 weeks. Asterisks indicate both GFP- and mCh-positive cells.

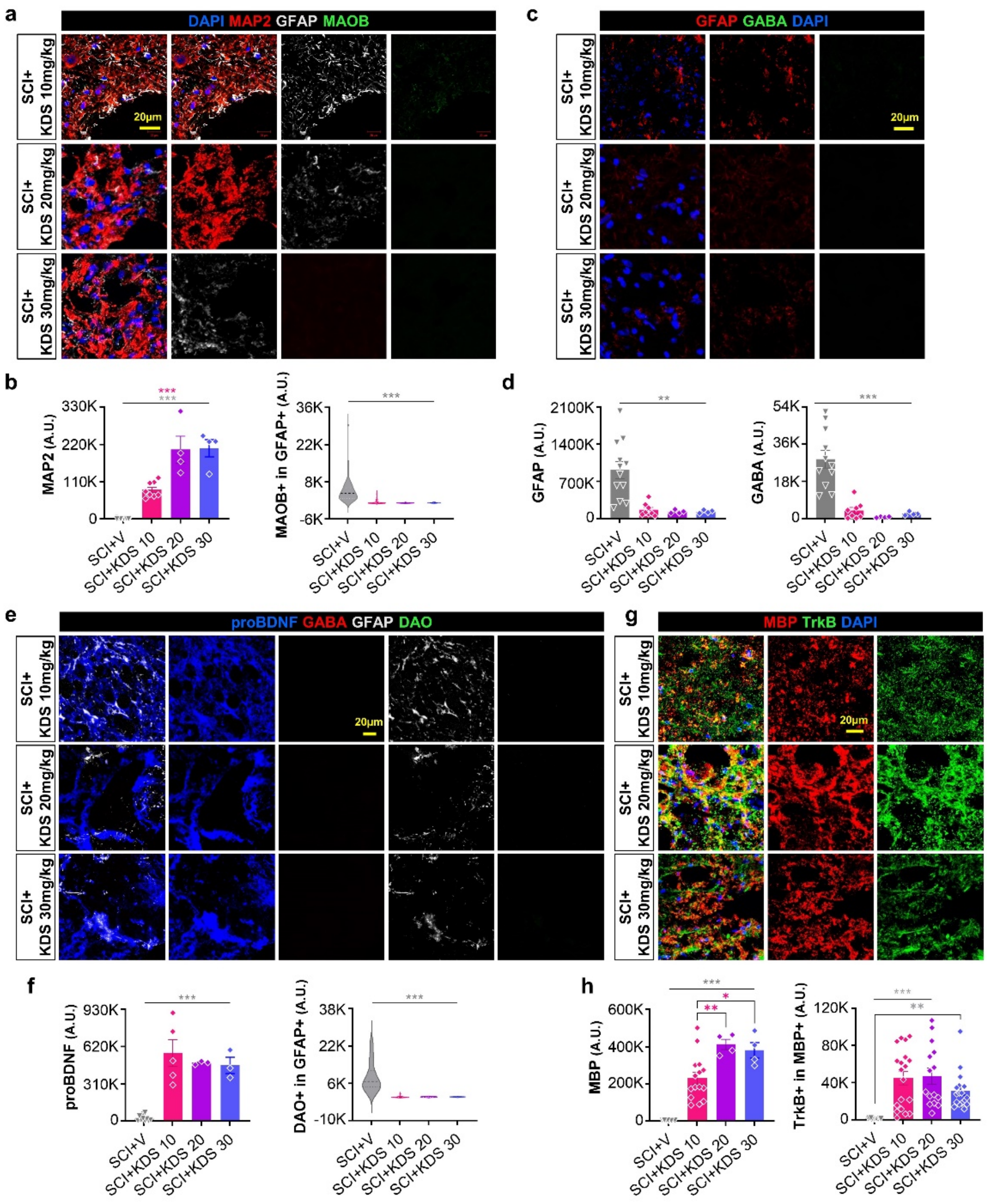


**Supplementary Fig. 6 KDS2010, at several concentrations, reduces astrocyte reactivity and GABA expression while enhancing proBDNF and TrkB expression.**

**a** Confocal images of the injured area in SCI+KDS 10mpk, 20mpk, and 30mpk stained with anti-MAP2 (red), GFAP (white), MAOB (green) antibodies, and DAPI (blue) at PI 10w. **b** (Left) The intensity of MAP2 showed a significant increase in SCI+KDS 10mpk, 20mpk, and 30mpk compared to SCI+V. All doses of the KDS induced a significant reduction in astrocytic MAOB expression (right) compared to SCI+V. **c** Confocal images of the injured area in SCI+KDS 10mpk, 20mpk, and 30mpk stained with anti-GFAP (red) and GABA (green) antibodies, and DAPI at PI 10w. **d** The intensities of GFAP (left) and GABA (right) in SCI+KDS 10mpk, 20mpk, and 30mpk were significantly reduced compared to SCI+V. **e** Confocal images of the injured area in SCI+KDS 10mpk, 20mpk, and 30mpk stained with anti-proBDNF (blue), GABA (red), GFAP (white), and DAO (green) antibodies at PI 10w. **f** The intensity of proBDNF significantly increased in SCI+KDS 10mpk, 20mpk, and 30mpk compared to SCI+V, while the intensity of astrocytic DAO significantly decreased. **g** Confocal images of the injured area in SCI+KDS 10mpk, 20mpk, and 30mpk stained with anti-MBP (red) and TrkB (green) antibodies, and DAPI (blue) at PI 10w. **h** (Left) The intensity of MBP showed a significant recovery in SCI+KDS 10mpk, 20mpk, and 30mpk compared to SCI+V. (Bottom) The intensity of MBP-positive TrkB showed a significant recovery in SCI+KDS 10mpk, 20mpk, and 30mpk compared to SCI+V.

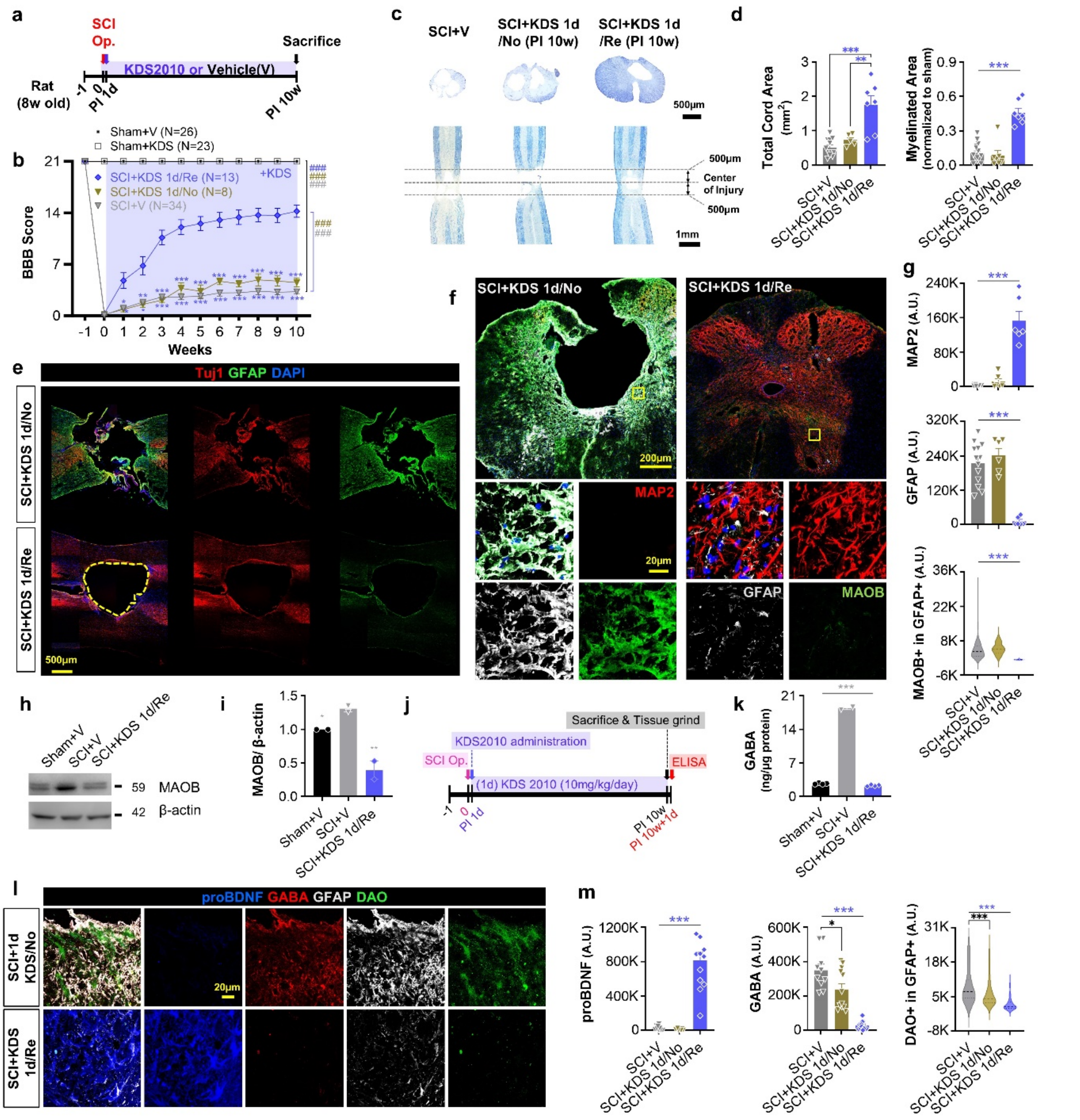


**Supplementary Fig. 7 MAOB inhibition from the acute phase induces recovery in some cases, while in others, recovery fails due to the compensatory action of DAO.**

**a** Experimental timelines using 8-week-old rats with the SCI operation and treatment with KDS2010, from 1 day after (acute) SCI. **b** SCI+KDS 1d/Re exhibited a stable gait at PI 10w, whereas SCI+KDS1d/No did not show any behavioral recovery in BBB locomotor test. Purple shade indicates the duration of KDS2010 administration. **c** EC staining of cross (top) and longitudinal (bottom) sections of spinal cord tissues in each group (SCI+V, SCI+KDS 1d/No, and SCI+KDS 1d/Re) at PI 10w. Dashed lines indicate region of analysis. **d** Reduced total spinal cord and myelinated areas in SCI+V and

SCI+KDS 1d/No were all significantly recovered in SCI+KDS 1d/Re at PI 10w. **e** Confocal images of longitudinal sections of spinal cord tissues stained with anti-Tuj1 (red), GFAP (green) antibodies, and DAPI (blue) at PI 10w in each group. **f** Confocal images of cross sections of spinal cord tissues in SCI+KDS 1d/No and SCI+KDS 1d/Re stained with anti-MAP2 (red), GFAP (white), MAOB (green) antibodies, and DAPI (blue) at PI 10w. **g** (Top) The intensity of MAP2 showed a significant increase in SCI+KDS 1d/Re compared to SCI+V and SCI+KDS 1d/No. In contrast, astrocytic GFAP (middle) and MAOB expression (bottom) significantly decreased in SCI+KDS 1d/Re compared to SCI+V and SCI+KDS 1d/No. **h** Western blotting of MAOB in Sham+V, SCI+V, and SCI+KDS 1d/Re at PI 10w. β-actin was used as a control for protein amount. **i** The MAOB protein level was significantly reduced in SCI+KDS 1d/Re compared to SCI+V. **j** Experimental timeline for ELISA using rats with SCI operation and acute phase KDS2010 treatment. **k** The concentration of GABA in Sham+V, SCI+V, and SCI+KDS 1d/Re at PI 10w. **l** Confocal images of the injured area in SCI+KDS 1d/No and SCI+KDS 1d/Re stained with anti- proBDNF (blue), GABA (red), GFAP (white), and DAO (green) antibodies. **m** (Left) The intensity of proBDNF was significantly increased in SCI+KDS 1d/Re compared to SCI+V and SCI+KDS 1d/No. The intensity of GABA (middle) and astrocytic DAO (right) were significantly reduced in SCI+KDS 1d/Re, while a significant level remained in SCI+V and SCI+KDS 1d/No.

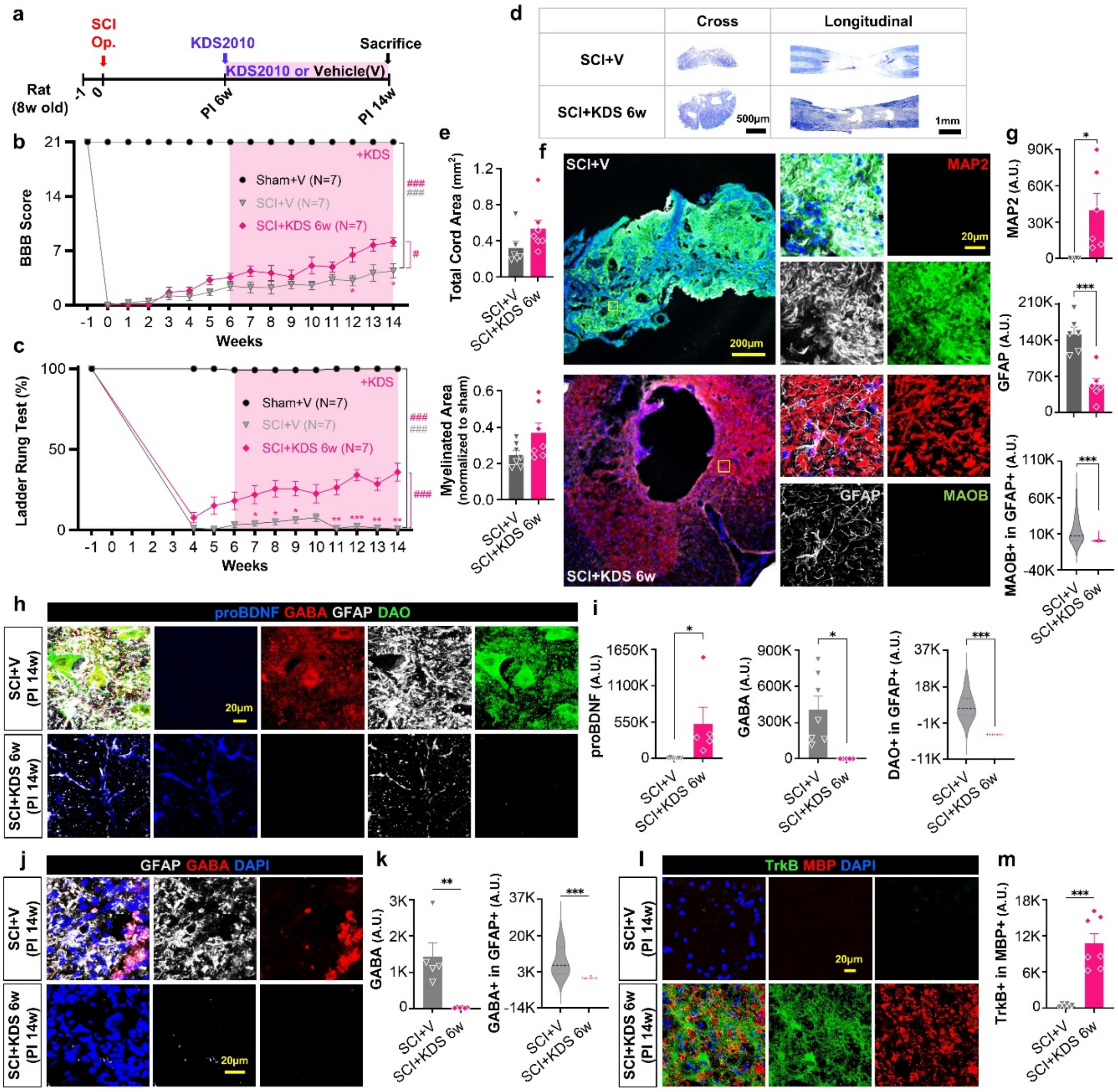


**Supplementary Fig. 8 MAOB inhibition from the chronic phase promotes recovery after SCI.**

**a** Experimental timelines using 8-week-old rats with the SCI operation and treatment with KDS2010, from 6 weeks after (chronic) SCI. **b, c** SCI+KDS 6w showed a significant motor recovery in BBB locomotor test (**b**) and ladder rung test (**c**). Pink shades indicate the duration of KDS2010 administration. **d** EC staining of cross (top) and longitudinal (bottom) sections of spinal cord tissues in SCI+V and SCI+KDS 6w at PI 14w. **e** The total spinal cord and myelinated areas in SCI+KDS 6w showed an increasing trend compared to SCI+V. **f** Confocal images of cross sections of spinal cord tissues in SCI+V and SCI+KDS 6w stained with anti-MAP2 (red), GFAP (white), MAOB (green) antibodies, and

DAPI (blue) at PI 14w. **g** (Top) The intensity of MAP2 showed a significant increase in SCI+KDS 6w compared to SCI+V. In contrast, astrocytic GFAP (middle) and MAOB expression (bottom) significantly decreased in SCI+KDS 6w compared to SCI+V. **h** Confocal images of the injured area in SCI+V and SCI+KDS 6w stained with anti- proBDNF (blue), GABA (red), GFAP (white), and DAO (green) antibodies. **i** (Left) The intensity of proBDNF was significantly increased in SCI+KDS 6w compared to SCI+V. The intensity of GABA (middle) and astrocytic DAO (right) were significantly reduced in SCI+KDS 6w, while a significant level remained in SCI+V and SCI+KDS 1d/No. **j** Confocal images of the injured area in SCI+V and SCI+KDS 6w stained with anti-GFAP (white), GABA (red) antibodies, and DAPI (blue) at PI 14w. **k** The elevated levels of astrocytic GFAP and GABA in SCI+V were significantly reduced in SCI+KDS 6w. **l** Confocal images of the injured area in SCI+V and SCI+KDS 6w stained with anti-TrkB (green) and MBP (red) antibodies, and DAPI (blue) at PI 14w. **m** The intensity of MBP-positive TrkB (bottom) was significantly increased in SCI+KDS 6w compared to SCI+V.

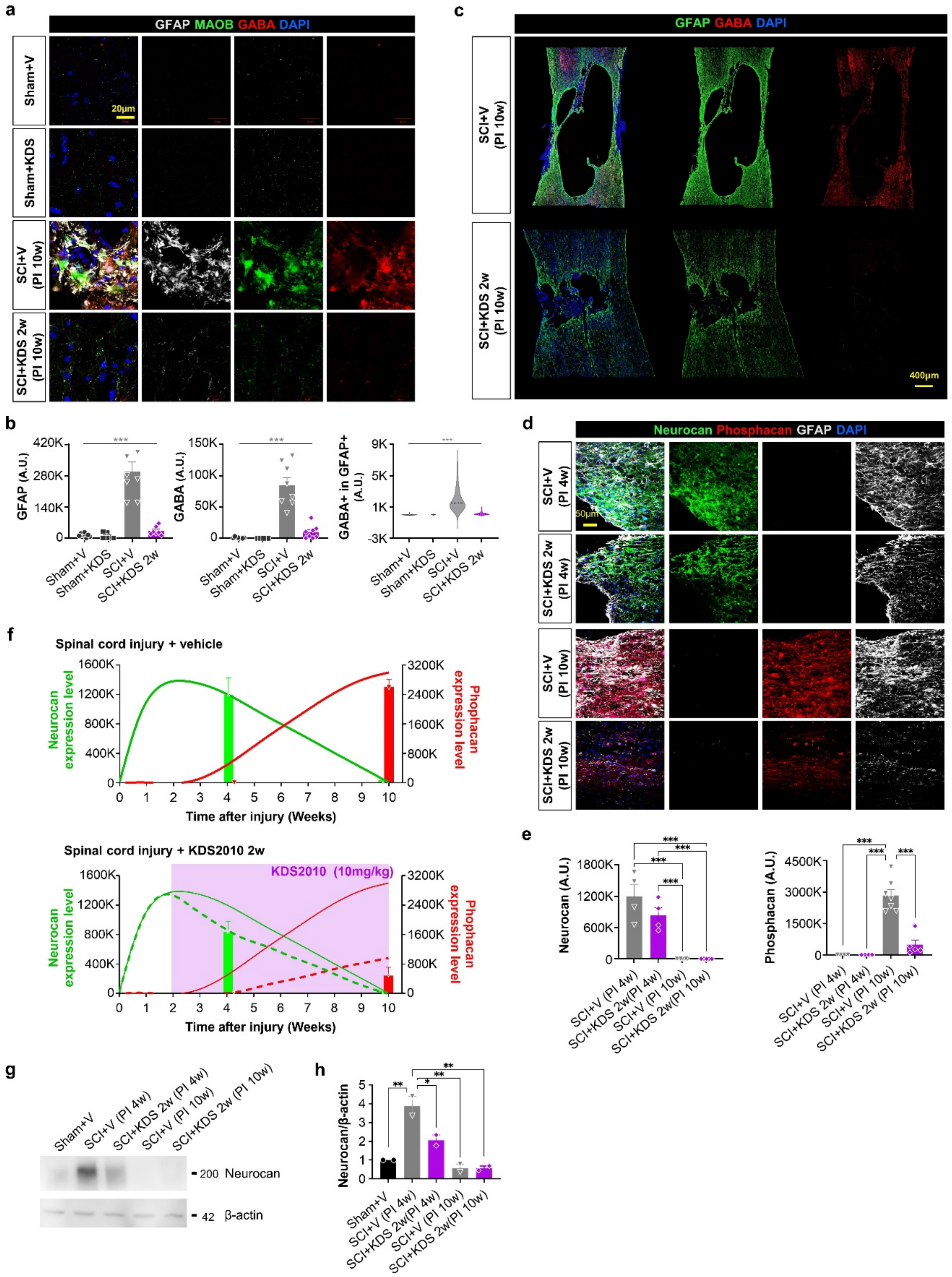


**Supplementary Fig. 9 MAOB inhibition reduces astrocytic GABA expression and CSPGs.**

**a** Confocal images of the injured areas in each group stained with anti-GFAP (white), MAOB (green), GABA (red) antibodies, and DAPI (blue) at PI 10w. **b** The intensity of GABA and astrocytic GABA, along with GFAP, significantly increased in SCI+V compared to Sham+V or Sham+KDS, while they significantly recovered in SCI+KDS 2w. **c** Confocal images of longitudinal sections of spinal cord tissues in SCI+V and SCI+KDS 2w stained with anti-GFAP (green), GABA (red) antibodies, and DAPI (blue) at PI 10w. **d** Confocal images of injured areas stained with anti-Neurocan (green), anti-Phosphacan (red), anti-GFAP (white), and DAPI at PI 4 and 10 weeks. **e** Quantification of Neurocan and Phosphacan intensities in each group. **f** Bar graph showing Neurocan and Phosphacan intensities over time, with dashed lines indicating expression trends in SCI+KDS 2w. **g** Western blot of Neurocan expression in Sham+V, SCI+V (PI 4w, 10w), and SCI+KDS 2w (PI 4w, 10w), normalized to β-actin. **h** Normalized Neurocan intensity relative to β-actin in each group.

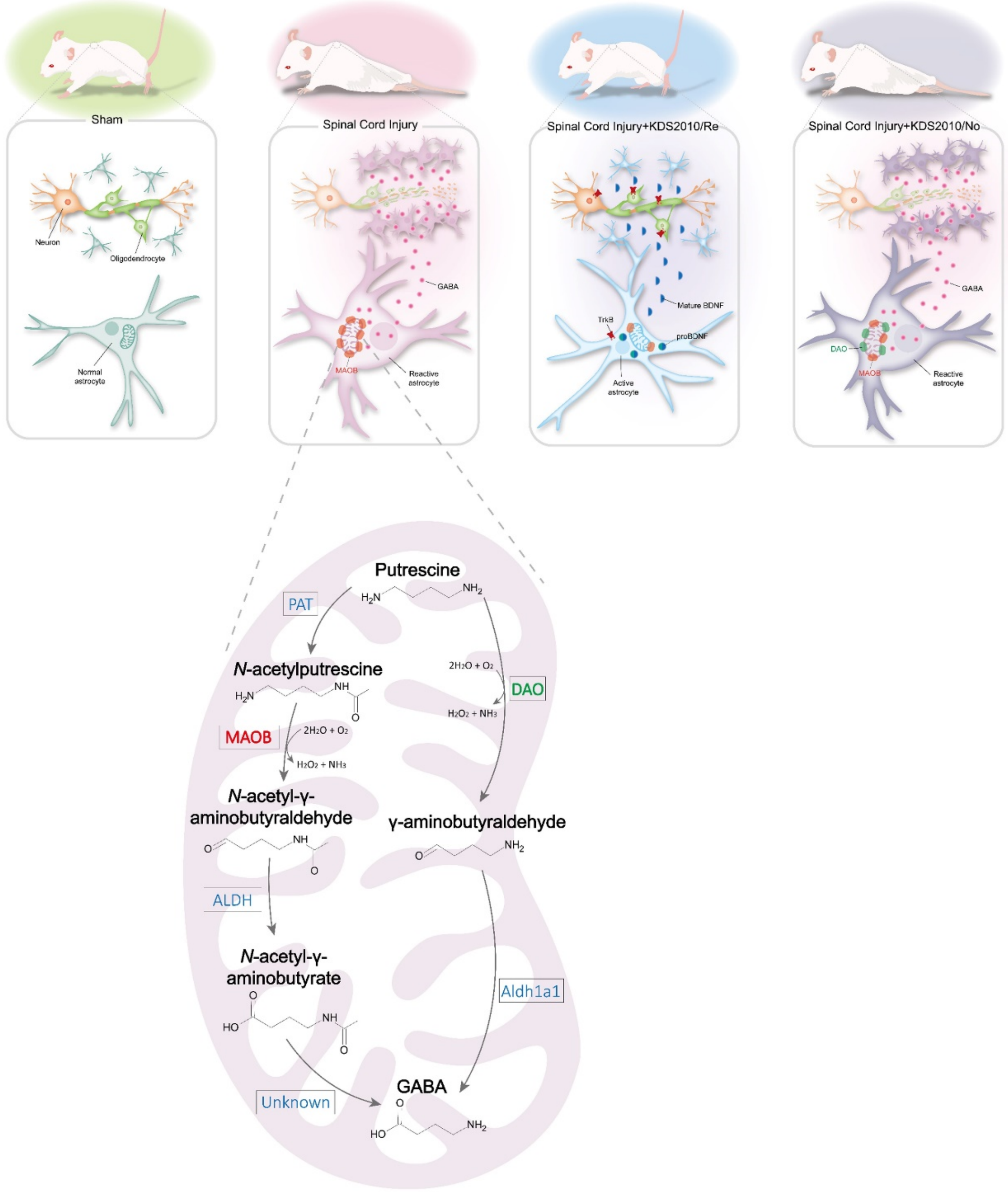


**Supplementary Fig. 10 Schematic model for molecular mechanism of neuroregeneration, remyelination, and functional recovery after SCI through MAOB inhibition.**

First, sham group possesses functional neurons, oligodendrocytes, and normal astrocytes. Second, SCI group shows reactive astrocytes with aberrant MAOB-dependent GABA production and release which impede neuroregeneration, remyelination, and functional recovery after SCI. Third, MAOB inhibition by KDS2010 leads to emergence of active astrocytes with increase of proBDNF and TrkB, and reduction of GABA. Finally, mBDNF, which is generated from cleavage of proBDNF, could cause neuroregeneration, remyelination, and functional recovery by acting on neuronal or oligodendrocytic TrkB which is also increased by MAOB inhibition in SCI+KDS2010/Re. Fourth, even if KDS2010 treated in acute phase, alternative pathway of DAO activated to degrade putrescine to GABA in reactive astrocytes, which consequently exerts a brake on proBDNF, neuroregeneration and functional recovery after SCI in SCI+KDS2010/No.

| | Single ascending dose study | | Multiple ascending dose study | | | |
|---|---|---|---|---|---|---|
| | Healthy young adults | | Healthy young adults | | Healthy elderly | |
| | Cohort 1-6 (n=36) | Placebo (n=12) | Cohort 7-10 (n=24) | Placebo (n=8) | Cohort 11 (n=6) | Placebo (n=2) |
| Age (years) | 29.33 ±5.74 | 28.50 ± 6.80 | 31.88 ± 6.10 | 31.50 ± 4.63 | 74.83 ± 2.32 | 71.50 ± 6.36 |
| Sex (n (%)) | | | | | | |
| Men | 35 (97.22) | 10 (83.33) | 24 (100.00) | 8 (100.00) | 6 (100.00) | 2 (100.00) |
| Women | 1 (2.78) | 2 (16.67) | 0 (0.00) | 0 (0.00) | 0 (0.00) | 0 (0.00) |
| Race, n (%) | | | | | | |
| Korean | 18 (50.00) | 7 (58.33) | 12 (50.00) | 4 (50.00) | 6 (100.00) | 2 (100.00) |
| Caucasian | 18 (50.00) | 5 (41.67) | 12 (50.00) | 4 (50.00) | 0 (0.00) | 0 (0.00) |
| Height (cm) | 175.35 ± 4.33 | 176.58 ± 7.28 | 174.95 ± 6.72 | 177.28 ± 8.20 | 164.10 ± 4.86 | 169.40 ± 0.99 |
| Weight (kg) | 74.34 ± 7.21 | 72.98 ± 7.89 | 73.48 ± 9.06 | 75.16 ± 8.51 | 62.58 ± 5.24 | 76.00 ± 5.94 |
| BMI (kg/m$^2$), | 24.20 ± 2.43 | 23.48 ± 2.97 | 23.98 ± 2.38 | 23.95 ± 2.56 | 23.23 ± 1.80 | 26.50 ± 2.40 |

**Supplementary Table 1 Demographic data in phase 1 clinical trial**

All data are presented as the mean ± standard deviation, unless otherwise specified.

| Cohort | N | $T_{max}$ ($T_{max,ss}$)(h) | $C_{max}$ ($C_{max,ss}$) (μg/L) | $AUC_{inf}$ ($AUC_{\tau,ss}$) (μg·h/L) | CL/F ($CL_{ss}$/F) (L/h) | Vd/F ($Vd_{ss}$/F) (L/h) | $t_{1/2}$ (h) | $CL_R$ (L/h) | Accumulation ratio* | Metabolic ratio** |
|---|---|---|---|---|---|---|---|---|---|---|
| **Single-ascending dose study** | | | | | | | | | | |
| Cohort 1 (30 mg) | 6 | 5.02 (0.93-8.00) | 43.26±9.91 | 2845.11±305.30 | 10.65±1.19 | 821.88 ± 80.51 | 53.93±7.21 | 0.07±0.02 | Not applicable | 11.06±3.83 |
| Cohort 2 (60 mg) | 6 | 3.00 (1.00-8.00) | 107.23±20.39 | 5546.66±927.35 | 11.02±1.48 | 678.87 ± 58.28 | 43.07±3.88 | 0.08±0.03 | | 11.66±3.22 |
| Cohort 3 (120 mg, fasting) | 6 | 3.00 (3.00-8.00) | 251.69±20.05 | 13850.72±2443.33 | 8.91±1.67 | 504.70 ± 76.10 | 40.19±8.01 | 0.07±0.04 | | 5.64±2.58 |
| Cohort 3 (120 mg, fed) | 5 | 6.00 (2.00-8.00) | 276.54±45.44 | 14283.46±3007.67 | 8.69±1.75 | 478.16 ± 64.75 | 39.19±7.89 | 0.07±0.04 | | 9.17±2.78 |
| Cohort 4 (240 mg) | 6 | 2.50 (2.00-6.00) | 504.82±43.26 | 27817.03±9388.05 | 9.31±2.52 | 575.74 ± 99.81 | 44.33±7.63 | 0.06±0.02 | | 6.89±3.63 |
| Cohort 5 (480 mg) | 6 | 7.00 (2.00-12.00) | 825.56±165.65 | 52664.56±11423.96 | 9.54±2.40 | 541.07 ± 85.01 | 40.48±8.11 | 0.07±0.03 | | 6.67±0.81 |
| Cohort 6 (960 mg) | 6 | 3.50 (2.00-6.00) | 2130.70±368.69 | 123140.68±13196.64 | 7.87±0.85 | 486.83 ± 69.81 | 42.89±4.70 | 0.05±0.02 | | 8.79±2.35 |
| **Multiple-ascending dose study, steady state** | | | | | | | | | | |
| Cohort 7 (60 mg) | 6 | 4.00 (3.00-6.00) | 367.10±33.65 | 7055.38±968.65 | 8.64±1.21 | 513.79 ± 18.34 | 45.06±7.84 | 0.21±0.05 | 3.35±0.50 | 7.19±2.85 |
| Cohort 8 (120 mg) | 6 | 3.50 (2.02-6.00) | 684.97±66.23 | 13382.80±1554.29 | 9.08±1.14 | 553.19 ± 102.71 | 47.08±9.42 | 0.20±0.06 | 3.30±0.49 | 7.69±2.75 |
| Cohort 9 (240 mg) | 6 | 3.00 (1.00-8.00) | 1715.62±273.28 | 32698.39±4795.90 | 7.47±1.10 | 445.69 ± 57.57 | 41.73±3.89 | 0.13±0.04 | 3.37±0.72 | 7.33±3.28 |
| Cohort 10 (480 mg) | 6 | 2.51 (1.00-6.00) | 2843.36±319.38 | 56268.23±5433.45 | 8.60±0.92 | 550.18 ± 101.01 | 41.50±5.74 | 0.12±0.02 | 3.02±0.39 | 7.30±2.35 |
| Cohort 11 (120 mg, elderly) | 6 | 3.50 (2.00-8.02) | 859.42±101.86 | 16749.55±1912.66 | 7.24±0.78 | 440.51 ± 28.69 | 50.32±5.44 | 0.20±0.06 | 3.03±0.25 | 4.69±3.32 |

**Supplementary Table 2 Pharmacokinetic parameters of KDS2010 in phase 1 clinical trial.**

All data are presented as the mean ± standard deviation, except $T_{max}$ ($T_{max,ss}$) which is presented as median (minimum – maximum). $C_{max}$, maximum concentration; $T_{max}$, time to reach the $C_{max}$; $AUC_{inf}$, area under the concentration-time curve from 0 to infinity; $AUC_{\tau,ss}$, area under the concentration-time curve during the dosing intervals at steady state; $t_{1/2}$, terminal half-life; CL/F, the apparent clearance; $CL_R$, renal clearance. *Accumulation ratio was calculated as the ratio of $AUC_{\tau,ss}$ after the administration of the seventh dose in comparison with $AUC_\tau$ after the administration of the first dose. **Metabolic ratio was calculated as the ratio of $AUC_{last,parent}$ in comparison with $AUC_{last,metabolite}$.

**Supplementary Movie 1 BBB score assessment of SCI+V and SCI+KDS 2w.**

(Left) SCI+V showed impaired motor function at PI 10w. (Right) SCI+KDS 2w showed a significant recovery of motor function at PI 10w.

**Supplementary Movie 2 Ladder rung test of SCI+V and SCI+KDS 2w.**

(Left) SCI+V exhibited severe motor dysfunction with hindlimb slipping during traversing the ladder at PI 10w. (Right) SCI+KDS 2w exhibited a significant recovery at PI 10w.
